# Major patterns in the introgression history of *Heliconius* butterflies

**DOI:** 10.1101/2023.06.21.545923

**Authors:** Yuttapong Thawornwattana, Fernando A. Seixas, Ziheng Yang, James Mallet

## Abstract

Gene flow between species is an important evolutionary process that can facilitate adaptation and lead to species diversification. It also makes reconstruction of species relationships difficult. Here, we use the full-likelihood multispecies coalescent (MSC) approach to estimate species phylogeny and major introgression events in *Heliconius* butterflies from whole-genome sequence data. We obtain a robust estimate of species branching order among major clades in the genus, including the “melpomene-silvaniform” group, which shows extensive historical and on-going gene flow. We obtain chromosome-level estimates of key parameters in the species phylogeny, including species divergence times, present-day and ancestral population sizes as well as the direction, timing, and intensity of gene flow. Our analysis leads to a phylogeny with introgression events that differ from those obtained in previous studies. We find that *H. aoede* most likely represents the earliest-branching lineage of the genus and that “silvaniform” species are paraphyletic within the melpomene-silvaniform group. Our phylogeny provides new, parsimonious histories for the origins of key traits in *Heliconius*, including pollen feeding and an inversion involved in wing pattern mimicry. Our results demonstrate the power and feasibility of the full-likelihood MSC approach for estimating species phylogeny and key population parameters despite extensive gene flow. The methods used here should be useful for analysis of other difficult species groups with high rates of introgression.

## Introduction

Introgression has been reported in a wide range of organisms, including humans and Neanderthals (Kuhlwilm et al. 2016), Darwin’s finches (Lamichhaney et al. 2018), felids (Li et al. 2016; Li et al. 2019), canids (Gopalakrishnan et al. 2018), horses (Jónsson et al. 2014; Gaunitz et al. 2018), living and extinct elephants (Palkopoulou et al. 2018), cichlid fishes (Malinsky et al. 2018), malaria mosquitoes (Fontaine et al. 2015; Small et al. 2020), *Drosophila* fruit flies (Suvorov et al. 2022), sunflowers (Rieseberg et al. 2003; Whitney et al. 2010), maize (Calfee et al. 2021), and yeast (Leducq et al. 2016). It is now widely recognized as an important process that can facilitate adaptation and speciation. While previous studies have confirmed the prevalence of gene flow, they give only a fragmented understanding of introgression because they mostly rely on approximate methods based on simple data summaries such as genome-wide site pattern counts or estimated gene trees. Those methods have limited power and cannot infer certain modes of gene flow or estimate most population parameters characterizing the process of species divergence and gene flow.

Recent advances make it possible to model genomic evolution under the multispecies coalescent (MSC) framework and use full-likelihood methods to estimate the species tree and quantify introgression (Wen and Nakhleh 2018; Zhang et al. 2018; Flouri et al. 2020). Analyses of both simulated and real data demonstrate that this full-likelihood MSC approach is efficient, accurate, and robust to moderate levels of model violation (Huang et al. 2022; Thawornwattana et al. 2023). A major advantage over approximate methods is the ability to estimate parameters of the species tree and introgression events among branches precisely, including the strength and direction of introgression as well as species divergence times, introgression times and effective population sizes. Current approximate methods are mostly unable to estimate these parameters or to infer gene flow between sister lineages (Jiao et al. 2021; Mirarab et al. 2021).

Neotropical butterflies of the genus *Heliconius* have become a model system for understanding introgression (Heliconius Genome Consortium 2012; Martin et al. 2013; Kozak et al. 2021; Cicconardi et al. 2023). Previous phylogenomic studies have demonstrated introgression among closely related species, but investigations of gene flow deeper among lineages were only partially successful: the various different methods yielded different phylogenies and introgression scenarios (Kozak et al. 2021; Cicconardi et al. 2023), perhaps due to methodological artefacts.

Recently, we demonstrated deep-level introgression and hybrid speciation in the erato-sara clade of *Heliconius* (Edelman et al. 2019; Thawornwattana et al. 2022). Here, we focus on the more complex melpomene crown group, or “melpomene-silvaniform” group, which includes the melpomene-cydno-timareta clade, and Brown’s (1981) “silvaniform” species that are mostly Müllerian co-mimics of Ithomiini models, together with the related *H. besckei* and *H. elevatus*. The melpomene-silvaniform species frequently hybridize today and laboratory crosses demonstrate some interfertility across the entire group (Mallet et al. 2007). Thus, extensive gene flow is likely, making estimation of the true species phylogeny difficult. We also examine the overall species divergence and deeper introgression history of the entire genus. By using full-likelihood analysis of whole-genome data, we overcome limitations of approximate methods to obtain a robust estimate of the species tree, with major introgression events between branches quantified in terms of direction, strength and timing. We use subsets of species to represent each major clade and to answer specific questions of introgression, which helps to keep the computation manageable. Our species phylogeny and introgression history provides a more parsimonious explanation for evolution of key traits in *Heliconius* than those previously inferred using approximate methods (Jay et al. 2018; Kozak et al. 2021; Cicconardi et al. 2023). We find evidence of extensive autosomal gene flow across the melpomene-silvaniform group, and show how trees based on the Z chromosome most likely represent the true species phylogeny. The so-called silvaniform species appear to be paraphyletic, contrary to previous findings based on approximate methods (Zhang et al. 2016; Kozak et al. 2021). As well as improving the understanding of diversification and gene flow in the genus *Heliconius*, we believe our approach provides useful pointers for studying species phylogeny with complex patterns of introgression in other taxa.

## Results

### *Ancestral gene flow at the base of* Heliconius *phylogeny*

We first establish phylogenetic relationships among six major clades of *Heliconius* using blocks of 200 loci (well-spaced short segments) across the genome, with each block spanning 1–3 Mb. We select a representative species from each major clade (*H. burneyi, H. doris, H. aoede, H. erato, H. sara*), three species from the melpomene-silvaniform group (*H. melpomene, H. besckei, H. numata*), and one outgroup species (*Eueides tales*) (**Tables S1–S3**). Species tree search in each block was carried out under the multispecies coalescent (MSC) model using the Bayesian program BPP (see **Methods**). Although each block was assumed to lack gene flow, different trees in each block across the genome are likely to result from introgression. Unlike a windowed concatenated tree approach, our MSC method takes into account incomplete lineage sorting and uncertainty in gene trees. Coding and noncoding loci were analyzed separately.

Two major patterns of species relationships were found in the genus *Heliconius*: scenario 1 (*erato-early*), with the erato-sara clade diverging first, is supported by ~75% of genomic blocks, while scenario 2 (*aoede-early*), with *H. aoede* diverging first, is supported by ~20% of blocks (**Figure 1A**; see **Table S4** for genome-level summaries, and **Table S5** for chromosome-level summaries). Using a different reference genome or more stringent filtering yielded similar fractions of the same species trees across the genome (**Figures S1** and **S2** and **Tables S4–S7**). In both scenarios, there is uncertainty concerning (i) the branching order of *H. doris* and *H. burneyi*, with *H. doris* diverging first being the most common relationship in both scenarios, and (ii) the branching order within the melpomene-silvaniform clade, with both *H. besckei* + *H. numata* and *H. melpomene* + *H. numata* being nearly equally common across the genome (**Figure 1B**,**C**). To focus on the deep divergences, we constructed simplified summaries of inferred species trees with *H. besckei, H. numata* and *H. melpomene* grouped together (“BNM” in **Figure 1A**; also see **Figure S2** and **Tables S4** and **S5**). Summaries of full species trees are in **Figure S1** and **Tables S6** and **S7**; posterior probabilities of these local blockwise trees are generally low, with the median probability for each MAP tree <0.7, reflecting both limited information from only 200 loci and the challenge of resolving short branches in the species trees.

**Figure 1.**
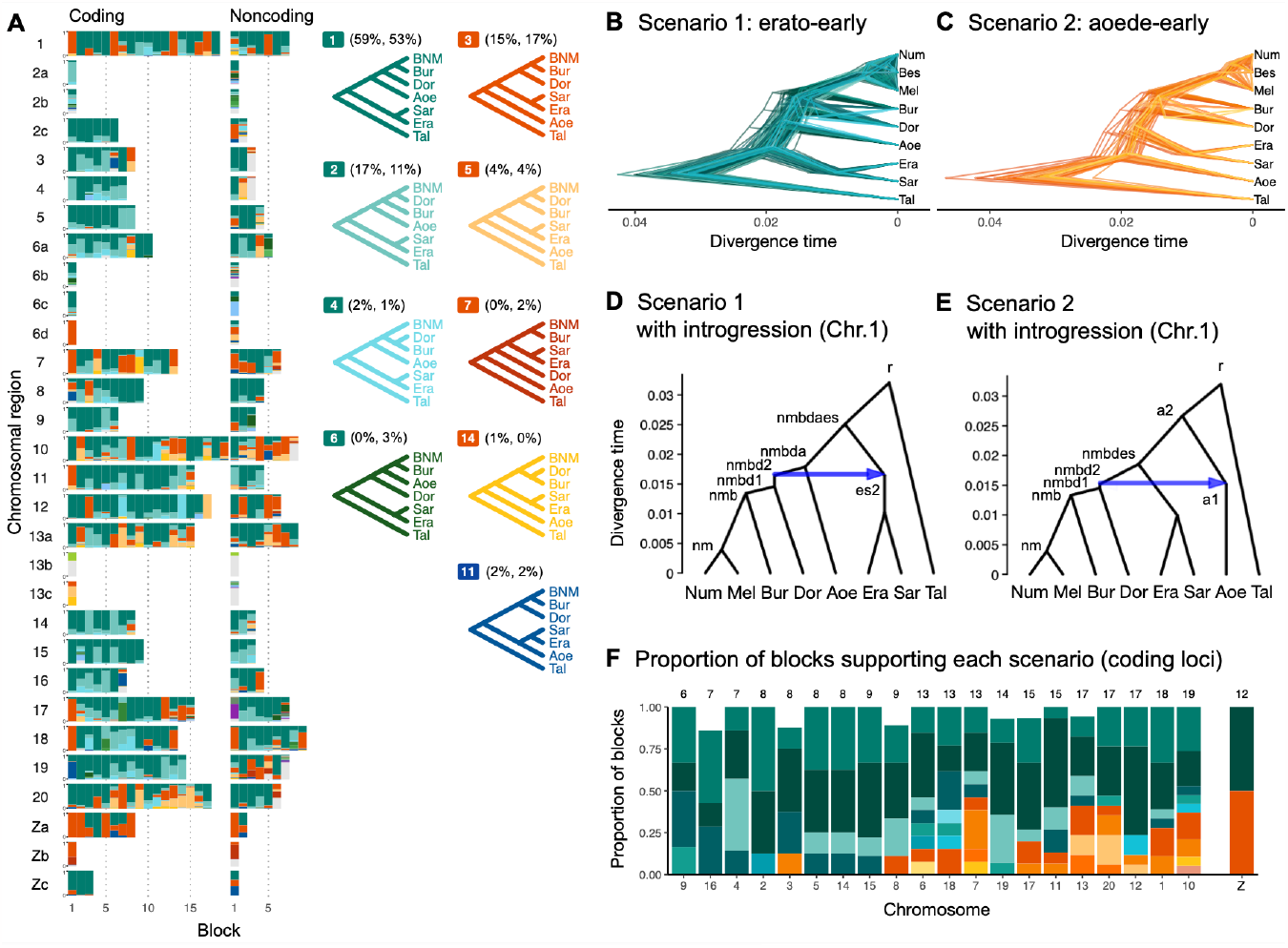
Ancestral gene flow at the base of *Heliconius*. (**A**) Posterior probabilities of species trees for major *Heliconius* clades inferred from BPP analysis of 200-locus blocks (each spanning 1–3 Mb) across the genome under the multispecies coalescent (MSC) model with no gene flow. The y-axis shows the posterior probability of species trees and ranges from 0 to 1. Colors correspond to the nine most common MAP trees, summarized by lumping species in the melpomene-silvaniform clade (Bes, Num, Mel) into a single tip (BNM); see **Figure S1** for full trees. Proportions of coding and noncoding blocks with each tree as a MAP tree are shown in parentheses. (**B, C**) Two scenarios of early divergence of *Heliconius*: (**B**) erato-early versus (**C**) aoede-early. Each tree is a MAP tree from a block having the MAP tree supporting one of these scenarios, with estimates of branch lengths (posterior means). (**D, E**) Phylogenetic and introgression histories estimated under an MSC model with introgression (MSC-I) corresponding to the two scenarios based on coding loci in chromosome 1 (3,517 coding loci; see **Figures S7** and **S8**). (**F**) Proportions of trees of scenarios 1 or 2 in each chromosome in order of increasing number of loci (used as a proxy for chromosome length). The Z chromosome (chr. 21) is placed at the right end. Number of blocks is shown on top of each bar. Tal: *Eueides tales*, Mel: *H. melpomene*, Bes: *H. besckei*, Num: *H. numata*, Bur: *H. burneyi*, Dor: *H. doris*, Aoe: *H. aoede*, Era: *H. erato*, Sar: *H. sara*.

Scenarios 1 and 2 are related via ancestral gene flow. If scenario 1 (**Figure 1B**) represents the true species tree, gene flow between the erato-sara clade and the common ancestor of *H. doris, H. burneyi* and the melpomene-silvaniform clade would lead to trees of scenario 2, with reduced estimated divergence time of the erato-sara clade. Similarly, if scenario 2 (**Figure 1C**) is the true tree, gene flow between the aoede clade and the common ancestor of *H. doris, H. burneyi* and the melpomene-silvaniform clade would lead to trees of scenario 1, with reduced estimated divergence time of *H. aoede*. This expected reduction in divergence time as a result of introgression is not apparent in our species tree estimates, partly due to a short internal branch separating *H. aoede* and the erato-sara clade in both scenarios (**Figure 1B,C**). To assess which scenario fits the data better, we calculated the Bayes factor under the MSC model with introgression (MSC-I) implemented in BPP, with *H. besckei* excluded to simplify the model (**Figure 1D,E**; ‘etales-8spp’ dataset in **Table S1**). The Bayes factor provides mixed evidence, with different chromosomes either strongly supporting alternative scenarios or not significant at 1% level; however the Z chromosome supports scenario 2 (**Table S11**). In scenario 2, divergence of *H. aoede* was estimated to be older than that of the erato-sara clade in scenario 1 while root age estimates were comparable, further supporting the hypothesis that *H. aoede* most likely diverged before the erato-sara clade (**Figure S9, Tables S9** and **S10**). This is because younger divergence may be explained by introgression whereas older divergence more likely represents true time of divergence. In scenario 2, introgression from the doris-burneyi-melpomene clade was estimated as unidirectional into *H. aoede* with a high probability (~75%), occurring shortly before the divergence of *H. doris* (**Figures S6** and **S8** and **Table S10**). Surprisingly, under scenario 1 we also find strong unidirectional introgression into the erato-sara clade with a high introgression probability (~65%) (**Figures S6** and **S7** and **Table S9**).

Two additional pieces of evidence support scenario 2. First, species trees of scenario 2 are most common in the Z chromosome (chr. 21) and tend to be more common in longer chromosomes, which have lower recombination rates per base pair (**Figure 1F**). Conversely, species trees of scenario 1 are more common in shorter chromosomes, which have higher recombination rates. If we assume that regions of low recombination tend to exhibit less introgression, with the Z chromosome being most resistant to gene flow in *Heliconius* (**Figures S3** and **S4** and **Table S8**), as suggested by previous studies (Edelman et al. 2019; Martin et al. 2019), the association of species trees of scenario 2 with regions of low recombination suggest that scenario 2 more likely represents the true species relationships whereas scenario 1 is a result of introgression. Second, we obtain a star tree whenever *H. erato* is assumed to diverge before *H. aoede* (scenario 1) under the MSC-with-migration (MSC-M or isolation-with-migration; IM) model of three species that allows for gene flow between two ingroup species since their divergence but does not allow gene flow with the outgroup species (**Figure S5**). Star trees are commonly observed when the assumed bifurcating tree was incorrect (Dalquen et al. 2017). By contrast, assuming *H. aoede* diverges before *H. erato* (scenario 2) always leads to a bifurcating tree.

Four small inversion regions (~100–400 kb) had been identified previously to be differentially fixed between the melpomene group and the erato-sara clade (Seixas et al. 2021): 2b, 6b, 6c, 13b and 21b. We were able to extract only a small number of loci (<100; see **Table S3**) from each region. While species tree estimates are more uncertain (**Figures S1** and **S2** and **Tables S4–S7**), the 13b region (~360 kb with respect to *H. erato demophoon* reference) consistently shows a unique pattern in which *H. doris* and *H. burneyi* cluster with the erato-sara clade instead of the melpomene-silvaniform clade. This suggests ancient introgression of an inversion from the erato-sara clade into *H. doris* and *H. burneyi* (Seixas et al. 2021).

In conclusion, we detect some hitherto unrecognized introgression events among the deepest branches within *Heliconius* sensu lato. The aoede-early scenario coupled with these deep introgression events may have led to some of the morphological and ecological evidence previously used in support of the erection the subgenera *Neruda* (for *H. aoede* and allies (Turner 1976)) as well as *Laparus* (for *H. doris* (Turner 1968)) that appeared to conflict with more recent molecular genetic data.

### Major introgression patterns in the melpomene-silvaniform clade

We next focus more closely on the melpomene-silvaniform clade (including *H. besckei, H. numata* and *H. melpomene*, represented by “BNM” in **Figure 1A**), one of the most phylogenetically difficult groups of *Heliconius* due to ongoing hybridization and extensive gene flow involving most members of the group (Mallet et al. 2007). Previous studies have inferred conflicting introgression scenarios in this clade (Martin et al. 2013; Zhang et al. 2016; Jay et al. 2018; Edelman et al. 2019; Cicconardi et al. 2023). We compiled a multilocus dataset from high-quality genome data comprising 8 (out of 15) species representing all major lineages within the clade: *H. melpomene, H. cydno, H. timareta, H. besckei, H. numata, H. hecale, H. elevatus* and *H. pardalinus* (**Table S12**). Our analysis below confirms widespread introgression within this clade.

We identify four major species relationships from blockwise estimates of species trees under the MSC model without gene flow (**Figures 2A** and **S10** and **Tables S13** and **S14**): (*a*) autosome-majority (trees 1–3), (*b*) autosome-variant (trees 5–7), (*c*) the Z chromosome (chr. 21; tree 4) and (*d*) chromosome 15 inversion region (15b; tree 24). The pattern is highly similar between coding and noncoding loci. The first three relationships (trees 1–9) account for >90% of the blocks, with a well-supported pardalinus-hecale clade ((*H. pardalinus, H. elevatus*), *H. hecale*), and a cydno-melpomene clade: ((*H. timareta, H. cydno*), *H. melpomene*). They differ in the position of *H. numata* and in the relationships among the three species in the cydno-melpomene clade. We first focus on the three scenarios relating to different positions of *H. numata*: (*a*) *H. numata* sister to the pardalinus-hecale clade + the cydno-melpomene clade, (*b*) *H. numata* sister to the pardalinus-hecale clade, and (*c*) *H. numata* sister to *H. besckei*. The Z-chromosome tree (i.e. tree 4 in scenario *c*) is the MAP tree with a high posterior probability in almost all blocks of the Z chromosome (median probability of 1; **Figure 2A** and **Table S13**), with *H. besckei* + *H. numata* diverging first, followed by a split between the pardalinus-hecale clade and the cydno-melpomene clade. A similar distinction between the autosome-majority trees and the Z-chromosome tree was also found by Zhang et al. (2016). All four scenarios *a*–*d* confirm paraphyly of the silvaniform species (*H. besckei, H. numata* and the pardalinus-hecale clade), consistent with some recent phylogenomic studies (Zhang et al. 2016; Massardo et al. 2020; Cicconardi et al. 2023). Monophyly of the silvaniforms was suggested in concatenation/sliding-window analysis (Heliconius Genome Consortium 2012; Kozak et al. 2015; Kozak et al. 2021; Zhang et al. 2021), but this conclusion may suffer from a failure to account for deep coalescence (Edwards et al. 2016). Our species tree search under the MSC (**Figures 1A** and **2A**) accounts for incomplete lineage sorting but does not account for gene flow.

**Figure 2.**
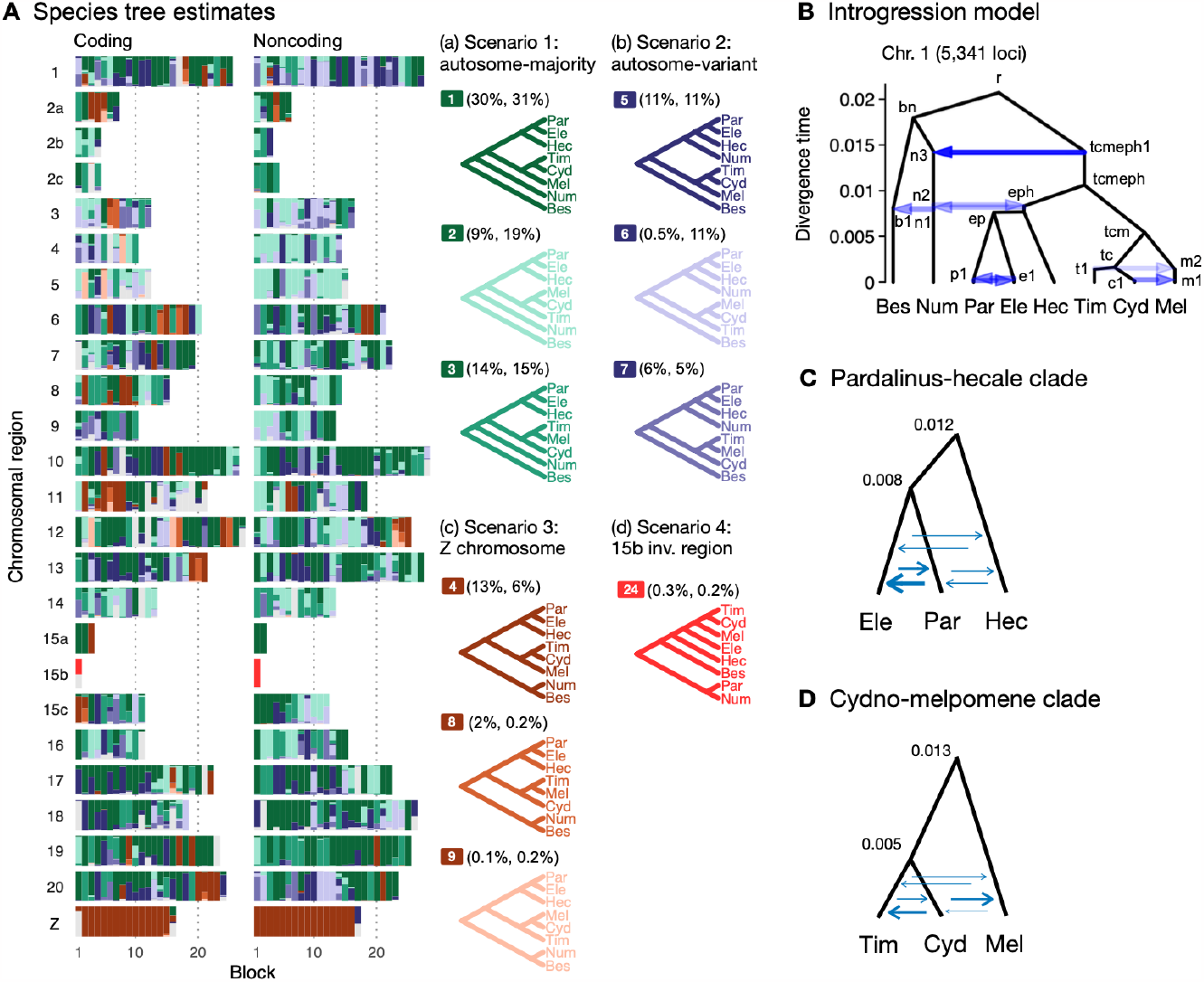
Major introgression events in the melpomene-silvaniform clade. (**A**) Blockwise estimates of species trees of the melpomene-silvaniform clade inferred from 200-locus blocks across the genome under the MSC model with no gene flow using BPP (see **Table S10** for data, **Tables S11** and **S12** and **Figure S10** for full results). MAP trees are labelled in decreasing order of frequency among blocks. Proportions of coding and noncoding blocks with each tree as a MAP tree are shown in parentheses. (**B**) The MSC-I model with introgression events that can explain the three major groups of trees in (**A**). Branch lengths are based on posterior means of divergence/introgression times estimated from 5,341 noncoding loci on chromosome 1 (**Table S15**). Each internal node is given a label, which is used to refer to a population above the node, e.g. the population between nodes r and bn is referred to as branch bn. Each horizontal arrow represents a unidirectional introgression event, e.g. the arrow from tcmeph1 to n3 represents tcmeph→Num introgression at time *τ*_tcmeph1_ = *τ*_n3_ with probability *φ*_tcmeph1_→_n3_. (**C**) Continuous migration (IM) model for the pardalinus-hecale clade, allowing bidirectional gene flow among the three species. (**D**) MSC-M model for the cydno-melpomene clade. For C and D, branch lengths are based on estimates from noncoding loci in chromosome 1 (**Figure S15**; **Tables S17–19**), and arrow sizes are proportional to estimated migration rate (*M* = *Nm*). Bes: *H. besckei*, Num: *H. numata*, Par: *H. pardalinus*, Ele: *H. elevatus*, Hec: *H. hecale*, Tim: *H. timareta*, Cyd: *H. cydno*, Mel: *H. melpomene*.

To approximate a fuller introgression history of this group, we construct a species tree model with introgression that can explain the four scenarios above. We use estimates of migration rates between each pair of species under the MSC-with-migration (MSC-M or IM) model of species triplets (3S analysis) to inform placement of introgression edges (**Figures S5** and **S11**). Our proposed model has six pairs of bidirectional introgression events (**Figure 2B**). We use the Z-chromosome tree (tree 4) as the backbone that most likely represents the true species tree. The other trees result from historical introgression. The tree is largely limited to the Z chromosome, which appears more resistant to gene flow in *Heliconius* (Zhang et al. 2016; Van Belleghem et al. 2018; Martin et al. 2019; Massardo et al. 2020; Thawornwattana et al. 2022). Consistent with this interpretation, we find no evidence of gene flow in the Z chromosome and high prevalence of gene flow on the autosomes based on the 3S analysis (**Figures S5** and **S11** and **Tables S15** and **S16**). Thus, we include two introgression events (between nodes n3–tcmeph1 and n2–eph1 in **Figure 2B**) to explain alternative positions of *H. numata* in the autosomes (scenarios *a* and *b*). Next, to explain the secondary source of genealogical variation within the cydno-melpomene clades (i.e. among trees 1–3, trees 4/8/9 and trees 5–7), we add two further introgression events between *H. melpomene* and *H. cydno* (m1–c1), and between *H. melpomene* and *H. timareta* (m2–t1). We do not model *H. cydno*–*H. timareta* introgression (these species are allopatric). We also include introgression between *H. besckei* and *H. numata* (b1–n1) to relate the autosome trees to the Z-chromosome tree. Finally, sister species *H. pardalinus* and *H. elevatus* hybridize today in sympatric populations (Rosser et al. 2019; Rosser et al. 2023), so we allow introgression between them (p1– e1). According to the 3S analysis, the rates of gene flow in this pair are among the highest (**Figure S11**). This sister-species introgression does not alter species trees (**Figure 2A)** because it does not change the topology.

The resultant MSC-with-introgression (MSC-I) model is used to estimate species divergence times, effective population sizes for extant and ancestral species, and the intensity, timing and direction of introgression. Consistent with scenario *c* representing the true species tree, we find least introgression on the Z chromosome (**Figures S12** and **S13** and **Tables S17** and **S18**). On the autosomes, there is substantial introgression from the pardalinus-hecale + cydno-melpomene clade into *H. numata*, and to a lesser extent, between *H. numata* and the pardalinus-hecale clade. These patterns match well with the frequencies of the two main autosomal relationships (scenarios *a* and *b* in **Figure 2**). Within the cydno-melpomene clade, introgression is predominantly unidirectional from *H. cydno* and *H. timareta* into *H. melpomene*. The *H. pardalinus*–*H. elevatus* pair shows on-going extensive but variable introgression across the genome, with the introgression time estimated to be zero. See SI text *‘Major introgression patterns in the melpomene-silvaniform clade inferred using 3s and BPP’* for more details.

The age of the melpomene-silvaniform clade (*τ*_r_) is estimated to be ~0.020 substitutions per site based on noncoding data (**Figure 2B** and **S13** and **Table S17**). This translates to ~1.7 (CI: 0.9, 3.8) million years ago (Ma), assuming a neutral mutation rate of 2.9×10^-9^ per site per generation (95% CI: 1.3×10^-9^, 5.5×10^-9^) and 4 generations per year (Keightley et al. 2015). This is not very different from a previous estimate of 3.7 (CI: 3.2, 4.3) Ma based on molecular clock dating (Kozak et al. 2015), which ignores ancestral polymorphism and is therefore expected to overestimate divergence time. Overall, our estimates of species divergence time tend to be precise and highly similar across the genome (**Figure S12**). The posterior means from coding and noncoding loci are strongly correlated, with *τ*_C_ ≈ *bτ*_NC_ where *b* varies between 0.4 and 0.6 (*r*^*2*^ > 0.95) in most chromosomal regions (**Figure S13**). The scale factor of *b* < 1 can be explained by purifying selection removing deleterious nonsonymous mutations in coding regions of the genome (Shi and Yang 2018). Present-day and ancestral population sizes (*θ*) are of the order of 0.01 (**Figure S13B** and **Tables S17** and **S18**). For inbred individuals (chosen for sequencing to facilitate genome assembly) among our genomic data (*H. melpomene, H. timareta, H. numata* and *H. pardalinus*; see **Table S1**), *θ* estimates vary among chromosomes by orders of magnitude, with the inbred genome of *H. melpomene* having the lowest population size of ~0.002–0.004 on average. Adding more individuals should help stabilize estimates of *θ*, but should not affect estimates of age or introgression rates.

The introgression model of **Figure 2B** assumes that gene flow occurs in single pulses. This may be unrealistic if gene flow is ongoing, so we also employ the MSC-M model implemented in BPP to estimate migration rates (*M* = *Nm*) between all pairs of species in each of the pardalinus-hecale and cydno-melpomene clades (**Figures 2C,D** and **S14** and **Table S19**). The MSC-M model assumes continuous gene flow since lineage divergence at the rate of *M* migrants per generation. The results concur with the introgression pulse model in suggesting high gene flow between *H. pardalinus* and *H. elevatus* (**Figure S15** and **Table S19**). There is also evidence of gene flow between *H. hecale* and *H. pardalinus*/*H. elevatus* at lower levels, with *M* < 0.1 in most chromosomes (**Figure S15** and **Table S19**). Allowing for continuous gene flow as well as gene flow involving *H. hecale*, we obtain slightly older estimates of both species divergence times (between *H. pardalinus* and *H. elevatus*, and between *H. hecale* and *H. pardalinus* + *H. elevatus*) (**Figure S15** and **Table S19**), than under the single-pulse introgression model (**Tables S17** and **S18**). The cydno-melpomene clade shows a similar pattern of older divergence times with substantial gene flow between all three species although at smaller magnitudes (*M* ~0.01–0.1) (**Table S21**). See SI text *‘MSC-M model for pardalinus-hecale and cydno-melpomene clades’* for more discussion of parameter estimates. We conclude that a model with continuous gene flow involving all three species may better describe the history of both the pardalinus-hecale clade and the cydno-melpomene clade.

In summary, we have identified substantial gene flow within the cydno-melpomene and pardalinus-hecale clades based on both pulse introgression (MSC-I) and continuous migration (MSC-M) models (**Figures 2D** and **S15, Tables S17–S21**). Earlier genomic studies failed to quantify the intensity of gene flow (introgression probability or migration rate) or infer direction and timing of gene flow. Gene flow within the cydno-melpomene clade has been extensively studied at population/subspecies levels and at specific loci involved in wing pattern mimicry (Bull et al. 2006; Pardo-Diaz et al. 2012; Kronforst et al. 2013; Martin et al. 2013; Wallbank et al. 2016; Enciso-Romero et al. 2017; Martin et al. 2019), but gene flow involving other species has received less attention (Heliconius Genome Consortium 2012; Wallbank et al. 2016; Zhang et al. 2016; Jay et al. 2018).

### Complex introgression in the 15b inversion region (P locus)

In the melpomene-silvaniform clade, a series of tandem inversions on chromosome 15 are involved in switching mimicry colour pattern in *H. numata* (Jay et al. 2018; Jay et al. 2022). The first inversion, P_1_ (~400 kb), is in the 15b region (also called the *P* locus), and is fixed in *H. pardalinus* and retained as a polymorphism in *H. numata* (Joron et al. 2011; Le Poul et al. 2014; Jay et al. 2018). Multiple introgression events are necessary to make the 15b tree (**Figure 2A**, scenario *d*, tree 24) compatible with either the Z chromosome tree or the autosomal trees (**Figure 2A**, scenarios *a*–*c*), suggesting a much more complex introgression history of this region than in the rest of the genome. This inversion contains the known wing patterning locus *cortex* (Jay et al. 2022), where it is maintained as a balanced polymorphism by natural selection (Joron et al. 2006; Nadeau et al. 2016; Van Belleghem et al. 2017). Another feature of this 15b region is that among the species without the inversion, the cydno-melpomene clade clusters with *H. elevatus* and is nested within the pardalinus-hecale clade (without *H. pardalinus*). This is contrary to the expectation based on the topologies in the rest of the genome (**Figure 2A**, scenarios *a*–*c*) that the cydno-melpomene clade would be sister to the pardalinus-hecale clade (without *H. pardalinus*). One explanation for this pattern is that introgression occurred between the common ancestor of the cydno-melpomene clade and either *H. elevatus* or the common ancestor of *H. elevatus* and *H. pardalinus* together with a total replacement of the non-inverted 15b in *H. pardalinus* by the P1 inversion from *H. numata* (Jay et al. 2018). We confirm and quantify this introgression below.

Using data from additional species (‘silv_chr15’ dataset in **Table S1**; see **Methods**), we obtain a better resolution of species relationships along chromosome 15, although with some uncertainty within the inversion region due to small numbers of loci (**Figure 3A** and **Table S22**). This analysis of independent data agrees with the Z chromosome tree (tree 24 in **Figure 2A**) and with (unrooted) trees obtained from concatenation analysis by Jay et al. (2018) (their Figures 2 and S1) and Jay et al. (2021) (their Figure S4), where *H. numata* with the inversion groups with *H. pardalinus* while *H. numata* without the inversion groups with its sister species, *H. ismenius* (**Figure 3A**, red trees). Outside the inversion region, *H. numata* with both inversion genotypes groups with *H. ismenius* as expected (**Figure 3A**, blue trees). Although this conclusion assumes that *H. numata* and *H. ismenius* are sister species while *H. ismenius* is not included in our species tree analysis of the melpomene-silvaniform clade (**Figure 2**), this sister relationship agrees with previous genomic studies of the autosomes and the sex chromosome (Zhang et al. 2016; Jay et al. 2021; Cicconardi et al. 2023; Rougemont et al. 2023).

**Figure 3.**
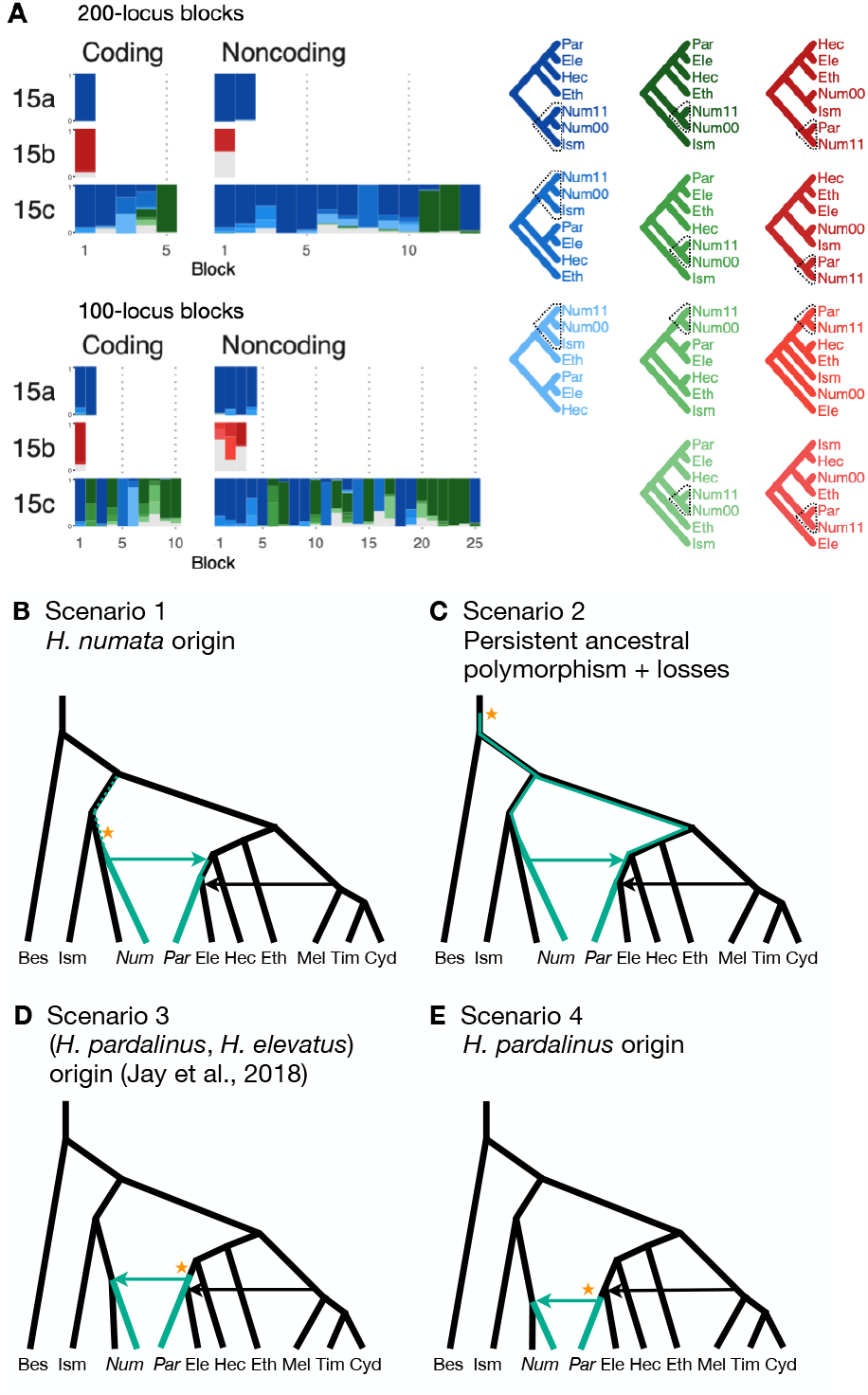
Introgression history of the chromosome 15 inversion region (15b) (**A**) Blockwise estimates of species trees for the inversion (15b) and the remnant flanking regions (15a and 15c) of chromosome 15. Trees were inferred from 200-locus blocks and 100-locus blocks under the MSC model without gene flow using BPP (**Table S20**). Tree legends are grouped by whether *H. numata* clusters with *H. ismenius* (blue), or *H. numata* with P_1_ inversion (Num11) clusters with *H. pardalinus* (red), or other relationships (green). (**B**–**E**) Four possible scenarios of the origin and introgression route of the P_1_ inversion. Star indicates the origin of the P_1_ inversion. Green lineages have the inversion, and green arrows indicates introgression of the inversion. Ism: *H. ismenius*, Num00: *H. numata* homozygous uninverted 15b (*H. n. laura* and *H. n. silvana*), Num11: *H. numata* homozygous for inversion P_1_ (*H. n. bicoloratus*), Eth: *H. ethilla*. See **Figure 2** legend for codes for other species. For the direction of melpomene-cydno clade→*H. elevatus*, see **Figure S18**.

Given that *H. numata* is an early-diverging lineage of the melpomene-silvaniform clade (**Figure 2B**) and is polymorphic for P_1_ over large parts of its geographic distribution while *H. pardalinus* is fixed for this inversion (Joron et al. 2011; Jay et al. 2022), one parsimonious explanation is that the inversion originated either in *H. numata* after diverging from *H. ismenius*, or before the *H. numata*– *H. ismenius* split but was subsequently lost in *H. ismenius*, followed by introgression which introduced the inversion from *H. numata* into *H. pardalinus*, either before or after its divergence from *H. elevatus* (**Figure 3B**). If the introgression occurred before *H. pardalinus*–*H. elevatus* divergence, the lack of the inversion in *H. elevatus* can be explained by another introgression from the cydno-melpomene clade into *H. elevatus*, completely replacing the inversion with the original orientation (**Figure 3B**). This introgression route is reported in previous genomic studies, including at a different locus (*optix* gene, also involved in colour pattern) in chromosome 18 as well as in the 15b region of chromosome 15 (Heliconius Genome Consortium 2012; Wallbank et al. 2016). Under this scenario, we might expect the 15b region to be less diverse in *H. pardalinus* (recipient of P_1_) than in *H. numata* (donor of P_1_), with the magnitude depending on the duration between the origin of P_1_ and introgression into *H. pardalinus*. However, we do not see this reduced heterozygosity in our data (**Figure S16**), suggesting that the transfer likely occurred early, shortly after the formation of the inversion in *H. numata*, or shortly after the *H. numata*–*H. ismenius* split if the inversion originated earlier. This scenario is further supported by the 15b tree having similar times of divergence between *H. ismenius* and *H. numata* without the inversion, and between *H. pardalinus* and *H. numata* with the inversion (**Figure S17**). The topology of the 15b tree (**Figures 2A** and **3A**) also indicates that the first split is between species with the inversion (*H. numata* and *H. pardalinus*) and those without the inversion. This suggests another possibility: inversion polymorphism could have existed earlier in the history of the melpomene-silvaniform clade but was subsequently lost in most species (**Figure 3C**). In this scenario, the ancestral polymorphism is maintained in *H. numata* while the inversion is fixed in *H. pardalinus* but is lost in other species. Introgression between *H. numata* and *H. pardalinus* is not required but could still occur.

To reconcile the introgression history of 15b with the overall species tree, we add three additional bidirectional introgression events onto the main model in **Figure 2B** and assess their plausibility using BPP. We allow for (i) bidirectional introgression between the cydno-melpomene clade into *H. elevatus*, (ii) bidirectional introgression between *H. numata* and *H. pardalinus*, and (iii) introgression between *H. besckei* and the common ancestor of the cydno-melpomene + pardalinus-hecale clade (to account for *H. besckei* being clustered with other species that do not have the inversion; see **Figure 2A**). We consider five models differring in the placements of introgression events (i) and (ii) either before or after the *H. pardalinus*–*H elevatus* split (**Figure S18**). Our results best support unidirectional introgression from *H. numata* into the common ancestor of *H. pardalinus* and *H. elevatus*, and from the common ancestor of the cydno-melpomene clade into *H. elevatus* shortly after its divergence from *H. pardalinus* (**Figure S18**, model m3). In other scenarios, estimated introgression times tend to collapse onto the *H. pardalinus*–*H elevatus* divergence time, suggesting that the introgression events were likely misplaced (**Figure S18** and **Table S23**). Our finding that divergence of *H. elevatus* and introgression from the cydno-melpomene clade occurred almost simultaneously provides evidence for a hybrid speciation origin of *H. elevatus* resulting from introgression between *H. pardalinus* and the common ancestor of the cydno-melpomene clade (Rosser et al. 2019; Rosser et al. 2023).

In an alternative scenario proposed by Jay et al. (2018), the inversion first originated in the common ancestor of *H. pardalinus* and *H elevatus*, and subsequently introgressed into some subspecies of *H. numata*, while the inversion in *H. elevatus* was completely replaced by introgression from the cydno-melpomene clade (**Figure 3D**). They used sliding-window gene tree topologies to support introgression of the inversion from *H. pardalinus* to *H. numata* shortly after its formation in the common ancestor of *H. pardalinus* and *H elevatus* (their Figures 4 and S4). By including *H. ismenius* and *H. elevatus*, sister species of *H. numata* and *H. pardalinus* respectively, different directions of introgression should lead to different gene tree topologies. Clustering of (*H. numata* with the inversion, *H. pardalinus*) with *H. numata* without the inversion would suggest the introgression is *H. numata* → *H. pardalinus* while clustering of (*H. numata* with the inversion, *H. pardalinus*) with *H. elevatus* would suggest *H. pardalinus* → *H. numata* introgression. However, tree topologies supporting each direction of introgression were almost equally common within the inversion region, particularly in the first half of the inversion, undermining this argument. With high levels of incomplete lineage sorting and introgression in the group, estimated gene trees need not reflect the true species relationships. Another variant of this scenario is that the inversion originated in *H. pardalinus* after its divergence from *H. elevatus*, and was introgressed into some subspecies of *H. numata* (**Figure 3E**). We consider this scenario unlikely because the inversion appears to originated long before the *H. pardalinus*–*H. elevatus* split given deep divergence of lineages with and without the inversion (see 15b trees in **Figures 2A** and **3A**).

**Figure 4.**
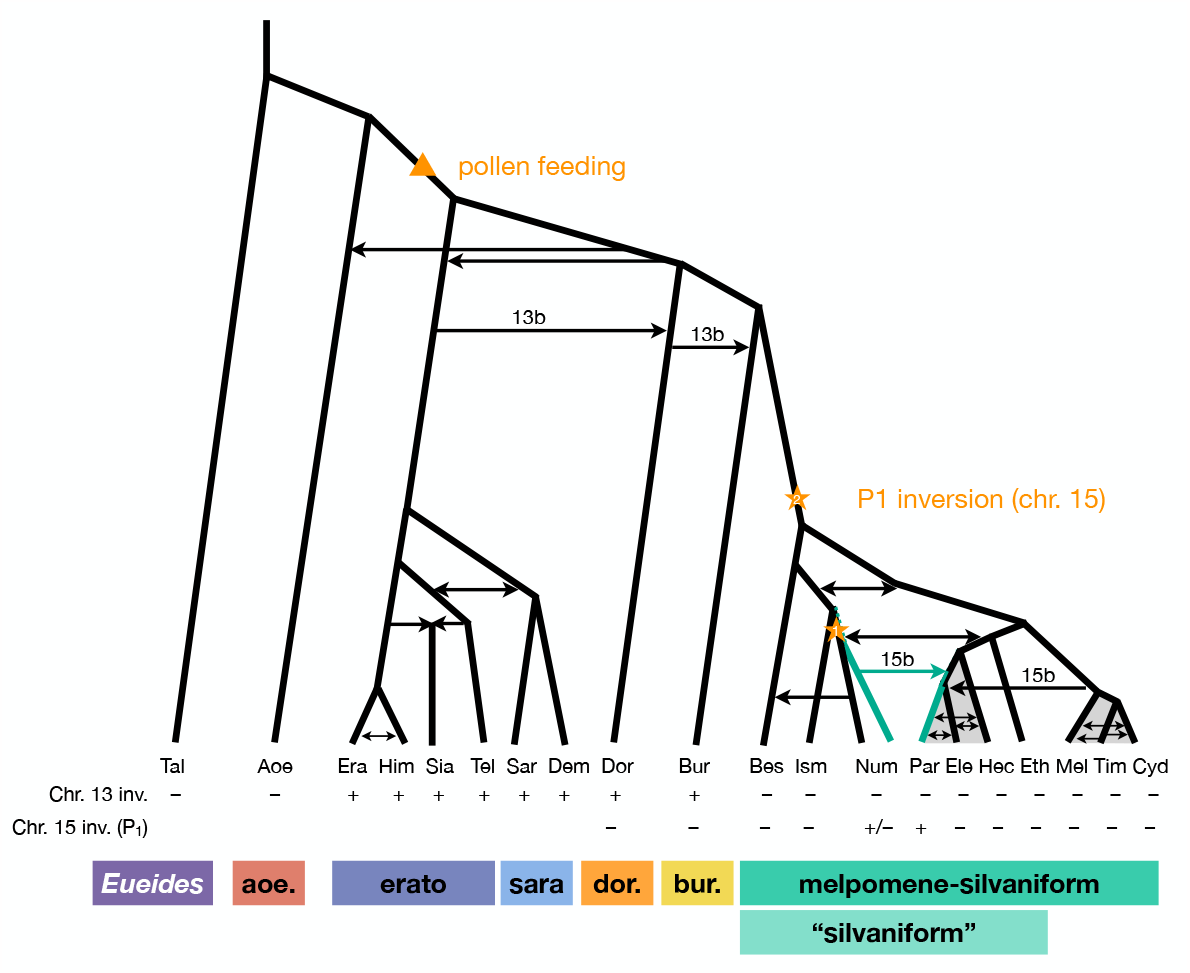
Revised phylogeny and introgression history of *Heliconius*. Phylogeny of the erato-sara clade was previously estimated using a similar approach (Thawornwattana et al. 2022). Arrows represent introgression events. Introgression of chromosomal inversions are region-specific, indicated by a label (13b or 15b) above the arrow. Status of inversions 13b and 15b in each species is indicated in the two rows at the bottom, with + indicating inversion, – indicating standard orientation, and +/– inversion polymorphism. Grey shading represents a period of continuous gene flow. Triangle represents the origin of pollen feeding near the base of *Heliconius*. Star indicates a possible origin of the P_1_ inversion on chromosome 15, and green branches indicate lineages with the inversion (polymorphic in Num, fixed in Par). Him: *H. himera*, Sia: *H. hecalesia*, Tel: *H. telesiphe*, Dem: *H. demeter*; see **Figures 1– 3** legends for other species codes.

Overall, we conclude that the P_1_ inversion likely arose in *H. numata* or its close ancestor and became fixed in *H. pardalinus* via an introgression replacement event from the former species (**Figure 3B**), although a deeper origin followed by persistent ancestral polymorphism (**Figure 3C**) remains a possibility.

## Discussion

### Approaches for estimating species phylogeny with introgression from whole-genome sequence data: advantages and limitations

Our full-likelihood MSC approach yields several improvements over concatenation and other approximate methods for inferring species trees and introgression. First, we account for incomplete lineage sorting (ILS) due to coalescent fluctuations, so variation in inferred genealogical histories can be more firmly attributed to variation in gene flow across the genome. Second, we analyze the sequence data directly, rather than using estimated gene trees as data. This utilizes information in branch lengths of the gene trees while accommodating their uncertainty. We retain heterozygous sites in the alignments as unphased diploid sequences from each individual without collapsing them into a single randomly-phased nucleotide as is common in other phylogenomic studies. Our approach allows multiple individuals per species to be included, leading to improved estimation of introgression parameters. Third, MSC models with pulse introgression (MSC-I) or continuous migration (MSC-M) allow direct estimation of key features of the direction, intensity, and timing of gene flow, and are applicable to gene flow between sister species. Widely-used summary statistics such as *D* and *f*_*d*_ do not estimate these parameters and cannot detect gene flow between sister lineages. Previous analyses using sliding-window concatenation or based on estimated gene trees were therefore liable to be less successful for estimating branching order in the melpomene-silvaniform clade, and because they may be misled by rapid speciation events coupled with extensive gene flow (Edelman et al. 2019; Massardo et al. 2020; Kozak et al. 2021).

There are several limitations in our approach. First, in the exploratory analysis of genomic blocks using MSC without gene flow (e.g. **Figures 1A** and **2A**), we assume that one of the main topologies represents the true species tree and is related to other topologies via introgression. In general, it can be nontrivial to decide which topology most likely represents the true species tree, and how to add introgression edges onto the assumed species tree to explain other topologies. Here, we employ several strategies including a recombination-rate argument (**Figure 1F**), reduced introgression in the sex chromosome (**Figure 2A**), estimates of pairwise migration rates under an explicit model of gene flow (**Figures S5** and **S11**), and using additional evidence from geographic distributions (Rosser et al. 2012) and records of natural hybrids (Mallet et al. 2007). These strategies could fail in taxa with more limited biological or distribution information, or when the sex chromosome (if available) is less unreliable as to species topology. Ideally, one would like to be able to estimate species phylogeny and introgression events simultaneously as part of a single unsupervised analysis, but this task remains computationally challenging. Software such as PhyloNet (Wen and Nakhleh 2018) and starBEAST (Zhang et al. 2018) implement Bayesian inference of MSC with introgression but they are usually limited to small datasets of no more than 100 loci. Heuristics based on summary statistics may attempt to place introgression edges onto a species phylogeny estimated on the assumption of no gene flow (Malinsky et al. 2018; Suvorov et al. 2022; Cicconardi et al. 2023). However, these approaches do not make full use of sequence data and can be misled by incorrect initial species trees. Furthermore, they cannot convincingly infer direction of gene flow or gene flow between sister species.

Second, although our approach is more powerful than approximate methods, it is also more computationally intensive and does not scale well to analyses of large numbers of species (>20, say). In *Heliconius*, there are 47 species in 6 major clades. Our strategy has therefore been to analyze subsets of species representing major clades or representative species within each clade. We then combine the results into a larger phylogeny. One caveat is that not all species get explored at the same time, and some introgression events may be missed. Lastly, the MSC makes simplifying modeling assumptions such as neutral evolution, constant substitution rates on all branches, no recombination within each locus, and free recombination among loci, the implications of which have been discussed previously (Burgess and Yang 2008; Thawornwattana et al. 2022). We note that although the MSC model assumes constant rate neutral evolution, estimated introgression probabilities (*φ*) are a combination of both actual introgression and subsequent positive and negative selection on introgressed loci, the extent of which is modulated by local recombination rate (Petry 1983; Barton and Bengtsson 1986).

In summary, we first explore major species relationships across genome blocks using a full-likelihood MSC approach. We then identify introgression events that explain the different local species trees by proposing introgression edges on a background species topology, with additional evidence from explicit models of gene flow. In addition, geographic distributions and the prevalence of natural hybrids can be employed to help with placement of introgression edges in a phylogeny. This process can result in several competing introgression models. We estimate parameters under each model using sequence data, evaluate the model fit, and perform model comparison. By breaking the problem into manageable parts in this way, our approach is computationally tractable.

### *An updated phylogeny of* Heliconius

We have clarified phylogenetic relationships among major clades within *Heliconius* and quantified major introgression events, including introgression between sister species (which does not change tree topology). We summarize our three key findings in **Figure 4**. First, *H. aoede* is most likely the earliest-branching lineage of *Heliconius*, and an ancient introgression event led to its apparently closer relationship with the doris-burneyi-melpomene clade (**Figure 1**). We discuss an implication of this placement below. However, we emphasize the considerable level of uncertainty that remains, and having more high-quality genome data from *H. aoede* and its sibling species may help improve the phylogenetic resolution. Second, within the melpomene-silvaniform clade, we obtained a robust pattern in which the common ancestor of *H. besckei, H. numata* and *H. ismenius* first split off from the rest, rendering the silvaniform species paraphyletic (**Figure 2**). This is contrary to a common belief that the silvaniforms are monophyletic (Heliconius Genome Consortium 2012; Kozak et al. 2015; Jay et al. 2021; Kozak et al. 2021; Zhang et al. 2021). Third, the P_1_ inversion on chromosome 15 involved in wing pattern mimicry likely introgressed from *H. numata* into *H. pardalinus* based on species tree topologies (**Figures 2** and **3**) and direct modeling of introgression (**Figure S18**), although where and when it originated remain uncertain.

*Heliconius* is an example of recent and rapid diversification with extensive gene flow occurring between closely related species as well as between more divergent lineages throughout its history at varying degrees. Since many species pairs of *Heliconius* show high degrees of sympatry today, there are plenty of opportunities for low levels of gene flow to occur whenever their ranges overlap (Rosser et al. 2015). This fact is evident in our estimates of pairwise migration rates among species in the melpomene-silvaniform clade (**Figures S3** and **S5**). There are many documented natural hybrids (Mallet et al. 2007) and there is evidence for introgression between particular pairs of subspecies from genomic studies, most of which involve wing patterning loci (Wallbank et al. 2016; Enciso-Romero et al. 2017; Morris et al. 2020; Rougemont et al. 2023). We did not attempt to incorporate all possible introgression signals into our phylogeny but instead focused on major introgression events that explain major patterns of species tree variation across the genome, using a representative subset of species. Thus our phylogeny in **Figure 4** should be viewed as a simplified version of reality. One future direction is to model a low level of admixture among all lineages.

### Implications for the evolution of pollen feeding

To illustrate how our updated phylogeny (**Figure 4**) can provide new insights into the evolution of *Heliconius*, we discuss an implication of our results for the evolution of pollen collection and pollen-feeding by adult *Heliconius*, a unique trait found in no other Lepidoptera (Gilbert 1972). Our phylogeny indicates that the aoede clade, comprising *H. aoede* and three other species, is likely the earliest-branching lineage of *Heliconius*, followed by the erato-sara clade. This branching order is consistent with a phylogeny based on morphological and behavioral characters (Penz 1999). Since the aoede clade is the only group of *Heliconius* that does not feed on pollen (Brown 1981), the aoede-early scenario (**Figure 1C**) suggests that unique pollen feeding traits arose once after the divergence of the aoede clade (**Figure 4**). In contrast, the erato-early scenario (**Figure 1B**) is less parsimonious as it requires an additional loss of pollen feeding ability in the aoede clade (Cicconardi et al. 2023).

Turner (1976) named the aoede clade as subgenus *Neruda* based on distinct morphology in all life stages, such as pupae that are more similar to those of *Eueides* (a sister genus of *Heliconius*), unique larva morphology, white eggs like those of *Eueides* (instead of yellow eggs in all other *Heliconius*), a different genus of larval host plant, *Dilkea* (Benson et al. 1975) (all other *Heliconius* species feed on *Passiflora*), and chromosome numbers of *n* = 21–31 which are variable and intermediate between *Heliconius* (*n* = 21 in most species) and *Eueides* (*n* = 31) (Brown et al. 1992). For these reasons, Brown (1981) later raised *Neruda* from subgenus to genus.

Thus, morphological/behavioral/ecological characters tend to support early divergence of the aoede clade (Brown 1972; Penz 1999). Molecular phylogenetic work, on the other hand, seemed to support the erato-early scenario (**Figure 1B**) (Brower and Egan 1997; Beltrán et al. 2007; Kozak et al. 2015; Kozak et al. 2021; Cicconardi et al. 2023). Consequently, generic status of *Neruda* was revoked (Kozak et al. 2015). Our full-likelihood approach, which accounts for incomplete lineage sorting as well as introgression, enables inference of a phylogeny that is more parsimonious with morphology and life history. This result parallels analysis of genomic data from African mosquitoes in the *Anopheles gambiae* species complex, in which coalescent-based likelihood analyses support species trees that are more parsimonious for chromosomal rearrangement data (Thawornwattana et al. 2018), while prior sliding-window and concatenation analysis favoured trees that are less parsimonious (Fontaine et al. 2015).

Our likelihood analyses can thus be used to rescue generic or subgeneric status for *Neruda*. Nevertheless, there remains considerable uncertainty near the base of *Heliconius*. Further whole-genome data from *H. aoede* and related species in the aoede, burneyi and doris clades will likely improve resolution.

## Methods

### Whole-genome sequence data and genotyping

We obtained raw sequencing data from previous studies (Wallbank et al. 2016; Edelman et al. 2019) (see **Table S1**) and extracted multilocus data as described previously (Thawornwattana et al. 2022) as unphased diploid sequences, retaining heterozygous sites. Sequencing reads were mapped to each of two reference genomes (see below) using bwa mem v0.7.15 (Li 2013) with default parameters, and then sorted using sambamba v0.6.8 (Tarasov et al. 2015). *RealignerTargetCreator* and *IndelRealigner* modules in GATK v3.8 were used to improve alignment around indels (McKenna et al. 2010; DePristo et al. 2011). We called genotypes of each individual using *mpileup* and *call* modules in bcftools v1.5 (Li et al. 2009) with the multiallelic-caller model (*call -m*) (Li 2011), and set minimum base and mapping quality to 20. We retained high-quality genotype calls using bcftools *filter* using the following filters: (1) a minimum quality score (QUAL) of 20 at both variant and invariant sites; (2) for each genotype, a genotype quality score (GQ) ≥ 20 and a read depth (DP) satisfying max(meanDP / 2, *d*) ≤ DP ≤ 2 meanDP, where *d* = 12 or 20 depending on the dataset, meanDP is the sample-averaged read depth. This choice of *d* was used to retain a large number of loci while maintaining low genotype-calling error rate (Thawornwattana et al. 2022). For the Z chromosome in females (which are heterogametic in *Heliconius*), we halved the DP threshold (*d*). Finally, we excluded sites within five base pairs (bp) of indels.

### Multilocus datasets

There are three main datasets in this study: (1) ‘etales-9spp’ contains eight species (*H. melpomene, H. besckei, H. numata, H. burneyi, H. doris, H. aoede, H. erato* and *H. sara*), one diploid individual per species, representative of all six major clades of *Heliconius* plus one species from a sister genus *Eueides* (*E. tales*). We used two reference genomes (*H. erato demophoon* v1 and *H. melpomene melpomene* Hmel2.5; see http://ensembl.lepbase.org/index.html) for read mapping, and two choices of the minimum read depth cutoffs (12 and 20) to ensure high-quality genotypes, resulting in four datasets in total. All reference genomes were available from lepbase.org. (2) ‘hmelv25-res’ contains eight species (one individual per species) within the melpomene-silvaniform clade (*H. melpomene, H. cydno, H. timareta, H. besckei, H. numata, H. hecale, H. elevatus* and *H. pardalinus*), mapped to Hmel2.5 reference. The DP threshold (*d*) for genotype calling was 20. (3) To understand the history of the 15b inversion region better, we also compiled a third multilocus dataset for chromosome 15 comprising the pardalinus-hecale clade, *H. numata* with and without the P_1_ inversion, as well as *H. ismenius* (sister species of *H. numata*) and *H. ethilla* (sister to the pardalinus-hecale clade) (**Table S1**, ‘silv_chr15’ dataset; see below under *‘Chromosome 15 inversion region: dataset and analysis’*) using publicly available data independent of our previous datasets used in this paper (Jay et al. 2018; Jay et al. 2021).

We generated coding and noncoding multilocus datasets from each dataset (**Table S1**) as follows (**Figure S20**). First, we extracted coordinates of coding and noncoding loci from the reference genome. In this study, loci refer to short segments of DNA that are far apart. The MSC model implemented in BPP assumes complete linkage within a locus and free recombination between loci. In simulations, species tree inference under MSC is found to be robust to within-locus recombination with recombination rates up to 10x the human rate (Zhu et al. 2022). Each coding locus coincided with a protein coding sequence (CDS) and had length at least 100 bp, whereas a noncoding locus can contain noncoding exons, introns and intergenic regions, and had length 100–1,000 bp. Since linkage disequilibrium in *Heliconius* species decays rapidly to background level within 10 kb (Heliconius Genome Consortium 2012), we spaced loci ≥2 kb apart, each assumed approximately independent, to obtain sufficient data. We excluded as loci any repetitive regions annotated in the reference genome. We processed each locus by removing sites containing missing data: the locus was discarded if it consisted of more than 50% missing data. After filtering, we also discarded loci with 10 or fewer sites remaining. We obtained >10,000 loci in each dataset; see **Tables S3** and **S12** for the number of loci. For the dataset with read depth cutoff of 12 aligned to the *H. erato* reference, we obtained about 19,000 noncoding loci and 48,000 coding loci. Note that we here obtain more coding than noncoding loci because noncoding loci were more difficult to align with the divergent *Eueides* outgroup (~7% divergent). Filtered noncoding loci were more conserved than coding loci for the same reason. The number of informative sites per locus was 10 for coding loci and 4 for noncoding loci on average. Average heterozygosity per site was about 0.43% for coding loci and 0.49% for noncoding loci, with *H. besckei* having the lowest heterozygosity (0.15–0.25%) and *H. burneyi* and *E. tales* having the highest heterozygosity (0.7–0.8%).

For the ‘etales-9spp’ dataset, we separated out inversion regions on chromosomes 2, 6, 13 and 21 into 2b, 6b, 6c, 13b and 21b (with two adjacent inversions in chromosome 6; chromosome 21 is the Z (sex) chromosome) while flanking regions were denoted 2a, 2c, 6a, 6d, 13a, 13c, 21a and 21c, resulting in 30 chromosomal regions in total. These inversions were first identified in a previous study (Seixas et al. 2021); see coordinates in **Table S2**. We obtained >11,000 noncoding loci and >31,000 coding loci in the smallest dataset (aligned to the *H. erato demophoon* reference, *d* = 20), and >31,000 noncoding loci and >48,000 coding loci in the largest dataset (Hmel2.5 reference, *d* = 12); see **Table S3**. The median number of sites was 100–130 depending on the dataset. The number of informative sites had median of 2–3 (range: 0–58) per locus for the noncoding loci and 4– 5 (0–570) for the coding loci. Again, noncoding loci are underrepresented in our datasets and they tended to be more conserved.

For the ‘hmvelv25-res’ dataset, we split chromosomes 2 and 15 into inversion (2b and 15b) and flanking regions (2a, 2c, 15a and 15c), resulting in 25 chromosomal regions in total; coordinates are in **Table S2**. Although only the chromosome 15b inversion region has been hitherto identified in this melpomene-silvaniform clade, we wished to test whether the chromosome 2b inversion identified in the erato-sara clade was also present in this group. We obtained >80,500 noncoding loci and >73,200 coding loci for ‘hmvelv25-res’ (**Table S12**). The median number of sites was 339 (range: 11–997) for noncoding loci and 147 (11–12,113) for coding loci. The median number of informative sites per locus was 6 (0–46) for noncoding loci and 2 (0–253) for coding loci.

### Overview of analysis approach

We first used the MSC model without gene flow to explore variation in genealogical history across the genome. We then formulated MSC models with introgression (MSC-I) based on a parsimony argument to explain major patterns of genealogical variation. We estimated the direction, timing, and intensity of introgression under each MSC-I model, and assessed most likely introgression scenarios. For gene flow between closely related species that may be on-going, we also used an MSC model with continuous migration (isolation-with-migration; IM) to estimate rates and directionality of gene flow.

### Species tree estimation under the MSC model without gene flow using BPP

We performed Bayesian inference of species trees under the MSC model without gene flow using BPP v4.4.0 (Yang and Rannala 2014; Rannala and Yang 2017; Flouri et al. 2018). This model accounts for gene-tree heterogeneity due to deep coalescence. Hence the genome-wide variation in estimated genealogy is most likely due to differential gene flow. We grouped loci into blocks of 200 and estimated a posterior distribution of species trees for each block. This blockwise analysis allows us to explore genealogical variation along each chromosomal region and to choose models of introgression for estimation in later analysis. Blocks of coding and noncoding loci were analyzed separately.

The MSC model without gene flow has two types of parameters: species divergence times (*τ*) and effective population sizes (*θ* = 4*Nμ*), both measured in expected number of mutations per site. For the ‘etales-9spp’ dataset (all four versions; **Figure 1** used H. erato reference depth greater than 12, “minDP12”), we assigned a diffuse gamma prior to the root age *τ*_0_ ~ G(7, 200), with mean 0.035, and to all population sizes *θ* ~ G(4, 200), with mean 0.02. Given *τ*_0_, other divergence times were assigned a uniform-Dirichlet distribution (Yang and Rannala 2010, eq. 2). For each block of loci, we performed ten independent runs of MCMC, each with 2×10^6^ iterations after a burn-in of 10^5^ iterations, with samples recorded every 200 iterations. We assessed convergence by comparing the posterior distribution of species trees among the independent runs. Non-convergent runs were discarded. The samples were then combined to produce the posterior summary such as the *maximum a posteriori* (MAP) tree. There were 1,355 blocks in total from all four versions of the dataset (**Table S3**), so there were 13,550 runs in total. Each run took about 20–30 hours (hrs).

Similarly, for the ‘hmelv25-res’ dataset, we used *τ*_0_ ~ G(4, 200), with mean 0.02, and population sizes *θ* ~ G(2, 200) for all populations, with mean 0.01. Each of the ten independent runs of the MCMC took 1×10^6^ iterations after a burn-in of 10^5^ iterations, with samples recorded every 100 iterations. There were 7,830 runs in total. Each run took about 15–20 hrs.

### Migration rate estimation under the MSC-M model for species triplets using 3S

To obtain more direct evidence of gene flow, we estimated migration rates between all pairs of *Heliconius* species in the ‘etales-8spp’ dataset under an MSC-with-migration (MSC-M or IM) model using the maximum likelihood program 3S v3.0 (Dalquen et al. 2017). The implementation in 3s assumes a species phylogeny of three species ((*S*_1_, *S*_2_), *S*_3_) with continuous gene flow between *S*_1_ and *S*_2_ since their divergence at constant rates in both directions, and requires three phased haploid sequences per locus. Since our multilocus data were unphased diploid, we phased the data using PHASE v2.1.1 (Stephens et al. 2001) to obtain two haploid sequences per individual at each locus. At each locus we then sampled three types of sequence triplets 123, 113 and 223 with probabilities 0.5, 0.25 and 0.25, respectively, where 113 means two sequences from *S*_1_ and one sequence from *S*_3_ chosen at random, etc. We used *E. tales* as the outgroup (*S*_3_) for all pairs (*S*_1_, *S*_2_). We analyzed coding and noncoding loci on the autosomes and three regions of the Z chromosome (chromosome 21) separately, each with 28 pairs among the eight *Heliconius* species. Additionally, we analyzed each autosomal region separately. This analysis was done with two choices of reference genome at read depth cutoff (*d*) of 12.

We fitted two models to each dataset: an MSC without gene flow (M0) and a bidirectional IM (M2). Model M0 has six parameters: two species divergence times (*τ*_1_ for *S*_1_–*S*_2_ divergence, *τ*_0_ for the root) and four population sizes (*θ*_1_ for *S*_1_, *θ*_2_ for *S*_2_, *θ*_4_ for the root, and *θ*_5_ for the ancestor of *S*_1_ and *S*_2_; there is no *θ*_3_ for *S*_3_ because there is at most one sequence from *S*_3_ per locus. Model M2 has two additional parameters: *M*_12_ and *M*_21_, where *M*_12_ = *m*_12_*N*_2_ is the expected number of migrants from *S*_1_ to *S*_2_ per generation, *m*_12_ is the proportion of migrants from *S*_1_ to *S*_2_ and *N*_2_ is the effective population size of *S*_2_. *M*_21_ is defined similarly. For each model, we performed ten independent runs of model fitting and chose the run with the highest log-likelihood. We discarded runs with extreme estimates (close to boundaries in the optimization). We then compared models M0 and M2 via likelihood ratio test (LRT), using a chi-squared distribution with two degrees of freedom as a null distribution at a significance threshold of 1%. Adjusting this threshold to account for multiple testing did not change our conclusions because the LRT values were usually extreme, especially in the analysis of all autosomal loci (**Table S8**). There were 31 (30 chromosomal regions + all of autosomal loci together) × 2 (coding and noncoding) × 28 (species pairs) × 2 (choices of reference genome) × 10 (replicates) = 34,720 runs in total. For our largest dataset with >46,000 loci, each run of fitting two models (M0 and M2) took 2–3 hrs.

We tested between two competing scenarios (**Figure 1B,C**) to infer which was more likely by estimating the internal branch length (Δ*τ* = *τ*_0_ – *τ*_1_) when either *H. aoede* or *H. erato* was used as an outgroup, with ingroup species representing the melpomene-silvaniform clade. To this end, we compiled another dataset similar to ‘hmelv25-res’ but included species from all six major clades of *Heliconius* (**Table S1**, ‘hmelv25-all’ dataset). We followed the same procedure as described above. There were 55 pairs in total for each choice of the outgroup species. Estimates of internal branch length close to zero (with the resulting tree becoming star-shaped) suggest that the specified species tree ((*S*_1_, *S*_2_), *S*_3_) was likely incorrect. There were 26 (25 chromosomal regions + all autosomal loci together) × 2 (coding and noncoding) × 55 (species pairs) × 2 (choices of reference genome) × 10 (replicates) = 57,200 runs in total.

### Parameter estimation under MSC with introgression (MSC-I) using BPP

Given the species-tree models with introgression of **Figure 1D,E**, we estimated introgression probabilities (*φ*), species divergence times and introgression times (*τ*) and effective population sizes (*θ*) for each coding and noncoding dataset from each chromosomal region using BPP v4.6.1 (Flouri et al. 2020). We assumed that population sizes of source and target populations remained unchanged before and after each introgression event (thetamodel = linked-msci). There were 25 unique parameters in total. We used the same prior distributions for *τ* and *θ* as in the MSC analysis without gene flow above, with root age *τ*_0_ ~ G(7, 200) and *θ* ~ G(4, 200) for all populations. We assigned a uniform prior U(0,1) to all introgression probabilities (*φ*). For each chromosomal region, we performed ten independent runs of MCMC, each with 1×10^6^ iterations after a burn-in of 10^5^ iterations, with samples recorded every 100 iterations. We assessed convergence by comparing the posterior estimates among the independent runs. Non-convergent runs were discarded. Samples were then combined to produce posterior summaries. Multiple posterior peaks, if they existed, were recorded and processed separately. There were 30 (chromosomal regions) × 2 (coding and noncoding) × 2 (trees 1 and 3) × 10 (replicates) = 1,200 runs in total. Each run took 200–300 hrs.

For the two MSC-I models of **Figure 1D,E**, we also estimated the marginal likelihood for each model using thermodynamic integration with 32 Gaussian quadrature points in BPP (Lartillot and Philippe 2006; Rannala and Yang 2017), and calculated Bayes factors to compare the two models (**Table S11**). This was done for each coding and noncoding dataset from each chromosomal region. Since the estimates of marginal likelihoods can be noisy, occasionally with extreme outliers, we adjusted estimates of Bayes factor by fitting local quadratic polynomials with span of 0.4 to the difference in the mean log likelihood from the two models, using the loess function in R (**Figure S19**). Additionally, we checked for reliability of the Bayes factor estimates by performing replicate calculation of the Bayes factor for a few chromosomes (3, 4, 9 and 21a) and found the preferred model choice to be reliable. The replicate estimates of log Bayes factor were -8.44, 19.35, -6.65, and 51.45 for chromosomes 3, 4, 9 and 21a, respectively, while the raw estimates were -11.41, 19.63, -10.94, and 53.12 and the adjusted estimates were -2.15, 20.83, -8.84, and 37.80 (**Table S11**).

The MSC-I model of **Figure 2B** has six pairs of bidirectional introgression events, with 12 introgression probabilities (*φ*), 13 species divergence times and introgression times (*τ*) and 15 population size parameters (*θ*), a total of 40 parameters. We assigned the same prior distributions to *τ*_0_ and *θ* as in the MSC analysis above, and *φ ~* U(0,1) for all introgression probabilities. The MSC-I model of **Figure 2C** has three more bidirectional introgression edges, with 9 additional parameters (6 introgression probabilities and 3 introgression times). Other settings were the same as above. There were 25 (chromosomal regions) × 2 (coding and noncoding) × 10 (replicates) = 500 runs in total. Most runs took 20–40 days.

### Parameter estimation under an MSC-M model using BPP

In the MSC-I model of **Figure 2B**, each of the pardalinus-hecale and cydno-melpomene clades has very recent estimated introgression times; thus gene flow may be on-going. We therefore used the MSC-M model to estimate six continuous migration rates between the three species in each clade using BPP v4.6.1 (Flouri et al. 2020), assuming the species tree as in **Figure 2B**. We assigned prior distributions *τ*_0_ ~ G(2, 200) for the root age (mean 0.01), *θ* ~ G(2, 500) for all populations (mean 0.004) and *M* ~ G(2, 10) for all migration rates (mean 0.2). The MCMC setup was the same as in the MSC-I analysis above. There were 25 (chromosomal regions) × 2 (coding and noncoding) × 10 (replicates) × 2 (pardalinus-hecale and cydno-melpomene clades) = 1,000 runs in total. Each run took 120–150 hrs.

### Chromosome 15 inversion region: dataset and analysis

To investigate the evolutionary history of the inversion region in chromosome 15 (15b region or *P* locus), we obtained genomic sequence data for six species (*H. numata* homozygous for the ancestral orientation, *H. numata* homozygous for the inversion, *H. ismenius, H. pardalinus, H. elevatus, H. hecale* and *H. ethilla*), with two individuals per species, from previous studies (Jay et al. 2018; Jay et al. 2021) (**Table S1**, ‘silv_chr15’ dataset). We extracted a multilocus dataset using an improved pipeline compared with that described above (see ‘*Whole-genome sequence data and genotyping’*, and ‘*Multilocus datasets’*). Here, a different filtering strategy used more complex, multi-stage genotyping to account for multiple individuals per species as follows. First, we removed Illumina adapters and trimmed low quality bases using trimmomatric v.0.39 (SLIDINGWINDOW:4:20 MINLEN:50). Next, sequencing reads were mapped to the *H. melpomene melpomene* reference assembly (Hmel2.5) using bwa-mem v.0.7.17 (Li 2013). Duplicate reads were masked using MarkDuplicates (Picard) in GATK v.4.2.6.1 (Poplin et al. 2018). We jointly called genotypes on chromosome 15 of individuals from the same species (and the same inversion genotype) using *mpileup* and *call* modules in bcftools v1.17 (Li et al. 2009) with the multiallelic-caller model (*call -m*) (Li 2011), and set the minimum base and mapping quality to 20. Only high-quality SNPs (QD score ≥2.0 and MQ score ≥40) were retained.

To obtain multilocus data, genomic coordinates of coding and noncoding loci were obtained from the reference genome as described above. Noncoding loci were 100 to 1000 bp in length and at least 2 kb apart. Coding loci were at least 100 bp in length (no maximum length limit) and at least 2 kb apart. To maximize information, minimum spacing was enforced for loci within the inversion region. We then extracted sequence alignments for each locus using the following procedure. All SNPs passing the quality filter were included. Constant sites were obtained from the reference sequence, unless they were masked by one of the following criteria: 1) read depth below 20 (coded as ‘–’), 2) non-SNP variant or low-quality SNP (coded as ‘N’). For each locus, we excluded sequences with >50% of sites missing (‘–’ or ‘N’), and excluded sites with all missing data. We discarded loci with only a single sequence remaining after filtering. Finally, we grouped loci into three regions as before: one inversion region (15b) and two flanking regions (15a and 15c). We obtained 218, 95, and 960 coding loci and 424, 368, and 2,446 noncoding loci in 15a, 15b, and 15c regions, respectively.

We performed blockwise estimation of species trees under the MSC model without gene flow as described earlier, in blocks of 100 and 200 loci (see ‘*Species tree estimation under the MSC model without gene flow using BPP’*). For the 200-locus blocks, there were 23 (7 coding blocks + 16 noncoding blocks) × 10 (replicates) = 230 runs; each run took about 60 hrs. For the 100-locus blocks, there were 45 (13 coding blocks + 32 noncoding blocks) × 10 (replicates) = 450 runs; each run took about 30 hrs.

## Supporting information

SI texts.

Figures S1–S20.

Tables S1–S23.

Available in Zenodo at https://doi.org/10.5281/zenodo.8415106.

## Data availability

All multilocus alignment datasets (see **Tables S1, S3** and **S12**) are available in Zenodo at https://doi.org/10.5281/zenodo.8415106.

## Acknowledgements

This study was supported by Harvard University (Y.T., F.S., and J.M.), and by Biotechnology and Biological Sciences Research Council grants (BB/T003502/1, BB/X007553/1 and BB/R01356X/1) and Natural Environment Research Council grant (NSFDEB-NERC NE/X002071/1) to Z.Y.

## References

Barton N, Bengtsson BO. 1986. The barrier to genetic exchange between hybridising populations. Heredity (Edinb). 57(3):357–376.

Van Belleghem SM, Baquero M, Papa R, Salazar C, McMillan WO, Counterman BA, Jiggins CD, Martin SH. 2018. Patterns of Z chromosome divergence among Heliconius species highlight the importance of historical demography. Mol Ecol. 27(19):3852–3872.

Van Belleghem SM, Rastas P, Papanicolaou A, Martin SH, Arias CF, Supple MA, Hanly JJ, Mallet J, Lewis JJ, Hines HM, et al. 2017. Complex modular architecture around a simple toolkit of wing pattern genes. Nat Ecol Evol. 1(3):52.

Beltrán M, Jiggins CD, Brower AVZ, Bermingham E, Mallet J. 2007. Do pollen feeding, pupal-mating and larval gregariousness have a single origin in Heliconius butterflies? Inferences from multilocus DNA sequence data. Biol J Linn Soc. 92(2):221–239.

Benson WW, Brown KS, Gilbert LE. 1975. Coevolution of Plants and Herbivores: Passion Flower Butterflies. Evolution (N Y). 29(4):659–680.

Brower AVZ, Egan MG. 1997. Cladistic analysis of Heliconius butterflies and relatives (Nymphalidae: Heliconiiti): a revised phylogenetic position for Eueides based on sequences from mtDNA and a nuclear gene. Proc R Soc London Ser B Biol Sci. 264(1384):969–977.

Brown KS. 1972. The heliconians of Brazil (Lepidoptera: Nymphalidae). Part III. Ecology and biology of Heliconius nattereri, a key primitive species neat extinction, and comments on the evolutionary development of Heliconius and Eueides. Zool New York. 57(2):41–69.

Brown KS. 1981. The Biology of Heliconius and Related Genera. Annu Rev Entomol. 26(1):427–457.

Brown KS, Emmel TC, Eliazar PJ, Suomalainen E. 1992. Evolutionary patterns in chromosome numbers in neotropical Lepidoptera: I. Chromosomes of the Heliconiini (Family Nymphalidae: Subfamily Nymphalinae). Hereditas. 117(2):109–125.

Bull V, Beltrán M, Jiggins CD, McMillan WO, Bermingham E, Mallet J. 2006. Polyphyly and gene flow between non-sibling Heliconius species. BMC Biol. 4(1):11.

Burgess R, Yang Z. 2008. Estimation of hominoid ancestral population sizes under Bayesian coalescent models incorporating mutation rate variation and sequencing errors. Mol Biol Evol. 25(9):1979–1994.

Calfee E, Gates D, Lorant A, Perkins MT, Coop G, Ross-Ibarra J. 2021. Selective sorting of ancestral introgression in maize and teosinte along an elevational cline. PLOS Genet. 17(10):e1009810.

Cicconardi F, Milanetti E, Pinheiro de Castro EC, Mazo-Vargas A, Van Belleghem SM, Ruggieri AA, Rastas P, Hanly J, Evans E, Jiggins CD, et al. 2023. Evolutionary dynamics of genome size and content during the adaptive radiation of Heliconiini butterflies. Nat Commun. 14(1):5620.

Dalquen DA, Zhu T, Yang Z. 2017. Maximum likelihood implementation of an isolation-with-migration model for three species. Syst Biol. 66(3):379–398.

DePristo MA, Banks E, Poplin R, Garimella K V., Maguire JR, Hartl C, Philippakis AA, Del Angel G, Rivas MA, Hanna M, et al. 2011. A framework for variation discovery and genotyping using next-generation DNA sequencing data. Nat Genet. 43(5):491–501.

Edelman NB, Frandsen PB, Miyagi M, Clavijo B, Davey J, Dikow RB, García-Accinelli G, Van Belleghem SM, Patterson N, Neafsey DE, et al. 2019. Genomic architecture and introgression shape a butterfly radiation. Science. 366(6465):594–599.

Edwards S V., Xi Z, Janke A, Faircloth BC, McCormack JE, Glenn TC, Zhong B, Wu S, Lemmon EM, Lemmon AR, et al. 2016. Implementing and testing the multispecies coalescent model: A valuable paradigm for phylogenomics. Mol Phylogenet Evol. 94:447–462.

Enciso-Romero J, Pardo-Díaz C, Martin SH, Arias CF, Linares M, McMillan WO, Jiggins CD, Salazar C. 2017. Evolution of novel mimicry rings facilitated by adaptive introgression in tropical butterflies. Mol Ecol. 26(19):5160–5172.

Flouri T, Jiao X, Rannala B, Yang Z. 2018. Species tree inference with BPP using genomic sequences and the multispecies coalescent. Mol Biol Evol. 35(10):2585–2593.

Flouri T, Jiao X, Rannala B, Yang Z. 2020. A bayesian implementation of the multispecies coalescent model with introgression for phylogenomic analysis. Mol Biol Evol. 37(4):1211–1223.

Fontaine MC, Pease JB, Steele A, Waterhouse RM, Neafsey DE, Sharakhov I V., Jiang X, Hall AB, Catteruccia F, Kakani E, et al. 2015. Extensive introgression in a malaria vector species complex revealed by phylogenomics. Science. 347(6217):1258524.

Gaunitz C, Fages A, Hanghøj K, Albrechtsen A, Khan N, Schubert M, Seguin-Orlando A, Owens IJ, Felkel S, Bignon-Lau O, et al. 2018. Ancient genomes revisit the ancestry of domestic and Przewalski’s horses. Science. 360(6384):111–114.

Gilbert LE. 1972. Pollen Feeding and Reproductive Biology of Heliconius Butterflies. Proc Natl Acad Sci. 69(6):1403–1407.

Gopalakrishnan S, Sinding MHS, Ramos-Madrigal J, Niemann J, Samaniego Castruita JA, Vieira FG, Carøe C, Montero M de M, Kuderna L, Serres A, et al. 2018. Interspecific Gene Flow Shaped the Evolution of the Genus Canis. Curr Biol. 28(21):3441–3449.e5.

Heliconius Genome Consortium. 2012. Butterfly genome reveals promiscuous exchange of mimicry adaptations among species. Nature. 487(7405):94–98.

Huang J, Thawornwattana Y, Flouri T, Mallet J, Yang Z. 2022. Inference of Gene Flow between Species under Misspecified Models. Mol Biol Evol. 39(12):msac237.

Jay P, Chouteau M, Whibley A, Bastide H, Parrinello H, Llaurens V, Joron M. 2021. Mutation load at a mimicry supergene sheds new light on the evolution of inversion polymorphisms. Nat Genet. 53(3):288–293.

Jay P, Leroy M, Le Poul Y, Whibley A, Arias M, Chouteau M, Joron M. 2022. Association mapping of colour variation in a butterfly provides evidence that a supergene locks together a cluster of adaptive loci. Philos Trans R Soc B Biol Sci. 377(1856):20210193.

Jay P, Whibley A, Frézal L, Rodríguez de Cara MÁ, Nowell RW, Mallet J, Dasmahapatra KK, Joron M. 2018. Supergene Evolution Triggered by the Introgression of a Chromosomal Inversion. Curr Biol. 28(11):1839–1845.e3.

Jiao X, Flouri T, Yang Z. 2021. Multispecies coalescent and its applications to infer species phylogenies and cross-species gene flow. Natl Sci Rev. 8(12):wab127.

Jónsson H, Schubert M, Seguin-Orlando A, Ginolhac A, Petersen L, Fumagalli M, Albrechtsen A, Petersen B, Korneliussen TS, Vilstrup JT, et al. 2014. Speciation with gene flow in equids despite extensive chromosomal plasticity. Proc Natl Acad Sci. 111(52):18655–18660.

Joron M, Frezal L, Jones RT, Chamberlain NL, Lee SF, Haag CR, Whibley A, Becuwe M, Baxter SW, Ferguson L, et al. 2011. Chromosomal rearrangements maintain a polymorphic supergene controlling butterfly mimicry. Nature. 477(7363):203–206.

Joron M, Papa R, Beltrán M, Chamberlain N, Mavárez J, Baxter S, Abanto M, Bermingham E, Humphray SJ, Rogers J, et al. 2006. A conserved supergene locus controls colour pattern diversity in Heliconius butterflies. PLoS Biol. 4(10):1831–1840.

Keightley PD, Pinharanda A, Ness RW, Simpson F, Dasmahapatra KK, Mallet J, Davey JW, Jiggins CD. 2015. Estimation of the Spontaneous Mutation Rate in Heliconius melpomene. Mol Biol Evol. 32(1):239–243.

Kozak KM, Joron M, McMillan WO, Jiggins CD. 2021. Rampant Genome-Wide Admixture across the Heliconius Radiation. Genome Biol Evol. 13(7).

Kozak KM, Wahlberg N, Neild AFE, Dasmahapatra KK, Mallet J, Jiggins CD. 2015. Multilocus species trees show the recent adaptive radiation of the mimetic heliconius butterflies. Syst Biol. 64(3):505–524.

Kronforst MR, Hansen MEB, Crawford NG, Gallant JR, Zhang W, Kulathinal RJ, Kapan DD, Mullen SP. 2013. Hybridization Reveals the Evolving Genomic Architecture of Speciation. Cell Rep. 5(3):666–677.

Kuhlwilm M, Gronau I, Hubisz MJ, De Filippo C, Prado-Martinez J, Kircher M, Fu Q, Burbano HA, Lalueza-Fox C, De La Rasilla M, et al. 2016. Ancient gene flow from early modern humans into Eastern Neanderthals. Nature. 530(7591):429–433.

Lamichhaney S, Han F, Webster MT, Andersson L, Grant BR, Grant PR. 2018. Rapid hybrid speciation in Darwin’s finches. Science. 359(6372):224–228.

Lartillot N, Philippe H. 2006. Computing Bayes factors using thermodynamic integration. Syst Biol. 55(2):195–207.

Leducq J-B, Nielly-Thibault L, Charron G, Eberlein C, Verta J-P, Samani P, Sylvester K, Hittinger CT, Bell G, Landry CR. 2016. Speciation driven by hybridization and chromosomal plasticity in a wild yeast. Nat Microbiol. 1(1):15003.

Li G, Davis BW, Eizirik E, Murphy WJ. 2016. Phylogenomic evidence for ancient hybridization in the genomes of living cats (Felidae). Genome Res . 26(1):1–11.

Li G, Figueiró H V., Eizirik E, Murphy WJ, Yoder A. 2019. Recombination-Aware Phylogenomics Reveals the Structured Genomic Landscape of Hybridizing Cat Species. Mol Biol Evol. 36(10):2111–2126.

Li H. 2011. A statistical framework for SNP calling, mutation discovery, association mapping and population genetical parameter estimation from sequencing data. Bioinformatics. 27(21):2987–2993.

Li H. 2013. Aligning sequence reads, clone sequences and assembly contigs with BWA-MEM. ArXiv e-prints.

Li H, Handsaker B, Wysoker A, Fennell T, Ruan J, Homer N, Marth G, Abecasis G, Durbin R. 2009. The Sequence Alignment/Map format and SAMtools. Bioinformatics. 25(16):2078–2079.

Malinsky M, Svardal H, Tyers AM, Miska EA, Genner MJ, Turner GF, Durbin R. 2018. Whole-genome sequences of Malawi cichlids reveal multiple radiations interconnected by gene flow. Nat Ecol Evol. 2(12):1940–1955.

Mallet J, Beltrán M, Neukirchen W, Linares M. 2007. Natural hybridization in heliconiine butterflies: The species boundary as a continuum. BMC Evol Biol. 7(1):28.

Martin SH, Dasmahapatra KK, Nadeau NJ, Salazar C, Walters JR, Simpson F, Blaxter M, Manica A, Mallet J, Jiggins CD. 2013. Genomewide evidence for speciation with gene flow in Heliconius butterflies. Genome Res. 23(11):1817–1828.

Martin SH, Davey JW, Salazar C, Jiggins CD. 2019. Recombination rate variation shapes barriers to introgression across butterfly genomes. PLoS Biol. 17(2):e2006288.

Massardo D, Vankuren NW, Nallu S, Ramos RR, Ribeiro PG, Silva-Brandão KL, Brandão MM, Lion MB, Freitas AVL, Cardoso MZ, et al. 2020. The roles of hybridization and habitat fragmentation in the evolution of Brazil’s enigmatic longwing butterflies, Heliconius nattereri and H. hermathena. BMC Biol. 18(1):84.

McKenna A, Hanna M, Banks E, Sivachenko A, Cibulskis K, Kernytsky A, Garimella K, Altshuler D, Gabriel S, Daly M, et al. 2010. The genome analysis toolkit: A MapReduce framework for analyzing next-generation DNA sequencing data. Genome Res. 20(9):1297–1303.

Mirarab S, Nakhleh L, Warnow T. 2021. Multispecies Coalescent: Theory and Applications in Phylogenetics. Annu Rev Ecol Evol Syst. 52(1):247–268.

Morris J, Hanly JJ, Martin SH, Van Belleghem SM, Salazar C, Jiggins CD, Dasmahapatra KK. 2020. Deep Convergence, Shared Ancestry, and Evolutionary Novelty in the Genetic Architecture of Heliconius Mimicry. Genetics. 216(3):765–780.

Nadeau NJ, Pardo-Diaz C, Whibley A, Supple MA, Saenko S V., Wallbank RWR, Wu GC, Maroja L, Ferguson L, Hanly JJ, et al. 2016. The gene cortex controls mimicry and crypsis in butterflies and moths. Nature. 534(7605):106–110.

Palkopoulou E, Lipson M, Mallick S, Nielsen S, Rohland N, Baleka S, Karpinski E, Ivancevic AM, To TH, Daniel Kortschak R, et al. 2018. A comprehensive genomic history of extinct and living elephants. Proc Natl Acad Sci U S A. 115(11):E2566–E2574.

Pardo-Diaz C, Salazar C, Baxter SW, Merot C, Figueiredo-Ready W, Joron M, McMillan WO, Jiggins CD. 2012. Adaptive introgression across species boundaries in Heliconius butterflies. PLoS Genet. 8(6):e1002752.

Penz CM. 1999. Higher level phylogeny for the passion-vine butterflies (Nymphalidae, Heliconiinae) based on early stage and adult morphology. Zool J Linn Soc. 127(3):277–344.

Petry D. 1983. The effect on neutral gene flow of selection at a linked locus. Theor Popul Biol. 23(3):300–313.

Poplin R, Ruano-Rubio V, DePristo MA, Fennell TJ, Carneiro MO, Van der Auwera GA, Kling DE, Gauthier LD, Levy-Moonshine A, Roazen D, et al. 2018. Scaling accurate genetic variant discovery to tens of thousands of samples. bioRxiv.:201178.

Le Poul Y, Whibley A, Chouteau M, Prunier F, Llaurens V, Joron M. 2014. Evolution of dominance mechanisms at a butterfly mimicry supergene. Nat Commun. 5(1):5644.

Rannala B, Yang Z. 2017. Efficient Bayesian species tree inference under the multispecies coalescent. Syst Biol. 66(5):823–842.

Rieseberg LH, Raymond O, Rosenthal DM, Lai Z, Livingstone K, Nakazato T, Durphy JL, Schwarzbach AE, Donovan LA, Lexer C. 2003. Major Ecological Transitions in Wild Sunflowers Facilitated by Hybridization. Science. 301(5637):1211–1216.

Rosser N, Kozak KM, Phillimore AB, Mallet J. 2015. Extensive range overlap between heliconiine sister species: evidence for sympatric speciation in butterflies? BMC Evol Biol. 15(1):125.

Rosser N, Phillimore AB, Huertas B, Willmott KR, Mallet J. 2012. Testing historical explanations for gradients in species richness in heliconiine butterflies of tropical America. Biol J Linn Soc. 105(3):479–497.

Rosser N, Queste LM, Cama B, Edelman NB, Mann F, Mori Pezo R, Morris J, Segami C, Velado P, Schulz S, et al. 2019. Geographic contrasts between pre- and postzygotic barriers are consistent with reinforcement in Heliconius butterflies. Evolution (N Y). 73(9):1821–1838.

Rosser N, Seixas FA, Queste LM, Cama B, Kryvokhyzha D, Mori-Pezo R, Nelson M, Waite-Hudson R, Goringe M, Costa M, et al. 2023. Hybrid speciation led to sympatric coexistence in Heliconius butterflies. Submitted.

Rougemont Q, Huber B, Martin SH, Whibley A, Estrada C, Solano D, Orpet R, McMillan WO, Frérot B, Joron M. 2023. Subtle Introgression Footprints at the End of the Speciation Continuum in a Clade of Heliconius Butterflies. Mol Biol Evol. 40(7):msad166.

Seixas FA, Edelman NB, Mallet J. 2021. Synteny-Based Genome Assembly for 16 Species of Heliconius Butterflies, and an Assessment of Structural Variation across the Genus. Genome Biol Evol. 13(7):evab069.

Shi CM, Yang Z. 2018. Coalescent-Based Analyses of Genomic Sequence Data Provide a Robust Resolution of Phylogenetic Relationships among Major Groups of Gibbons. Mol Biol Evol. 35(1):159–179.

Small ST, Labbé F, Lobo NF, Koekemoer LL, Sikaala CH, Neafsey DE, Hahn MW, Fontaine MC, Besansky NJ. 2020. Radiation with reticulation marks the origin of a major malaria vector. Proc Natl Acad Sci. 117(50):31583–31590.

Stephens M, Smith NJ, Donnelly P. 2001. A new statistical method for haplotype reconstruction from population data. Am J Hum Genet. 68(4):978–989.

Suvorov A, Kim BY, Wang J, Armstrong EE, Peede D, D’Agostino ERR, Price DK, Waddell PJ, Lang M, Courtier-Orgogozo V, et al. 2022. Widespread introgression across a phylogeny of 155 Drosophila genomes. Curr Biol. 32(1):111–123.e5.

Tarasov A, Vilella AJ, Cuppen E, Nijman IJ, Prins P. 2015. Sambamba: Fast processing of NGS alignment formats. Bioinformatics. 31(12):2032–2034.

Thawornwattana Y, Dalquen D, Yang Z. 2018. Coalescent analysis of phylogenomic data confidently resolves the species relationships in the Anopheles gambiae species complex. Mol Biol Evol. 35(10):2512–2527.

Thawornwattana Y, Huang J, Flouri T, Mallet J, Yang Z. 2023. Inferring the Direction of Introgression Using Genomic Sequence Data. Mol Biol Evol. 40(8):msad178.

Thawornwattana Y, Seixas FA, Yang Z, Mallet J. 2022. Full-Likelihood Genomic Analysis Clarifies a Complex History of Species Divergence and Introgression: The Example of the erato-sara Group of Heliconius Butterflies. Syst Biol. 71(5):1159–1177.

Turner JRG. 1968. Some new Heliconius pupae: their taxonomic and evolutionary significance in relation to mimicry (Lepidoptera, Nymphalidae)*. J Zool. 155(3):311–325.

Turner JRG. 1976. Adaptive radiation and convergence in subdivisions of the butterfly genus Heliconius (Lepidoptera: Nymphalidae). Zool J Linn Soc. 58(4):297–308.

Wallbank RWR, Baxter SW, Pardo-Diaz C, Hanly JJ, Martin SH, Mallet J, Dasmahapatra KK, Salazar C, Joron M, Nadeau N, et al. 2016. Evolutionary Novelty in a Butterfly Wing Pattern through Enhancer Shuffling. PLoS Biol. 14(1):e1002353.

Wen D, Nakhleh L. 2018. Coestimating reticulate phylogenies and gene trees from multilocus sequence data. Syst Biol. 67(3):439–457.

Whitney KD, Randell RA, Rieseberg LH. 2010. Adaptive introgression of abiotic tolerance traits in the sunflower Helianthus annuus. New Phytol. 187(1):230–239.

Yang Z, Rannala B. 2010. Bayesian species delimitation using multilocus sequence data. Proc Natl Acad Sci U S A. 107(20):9264–9269.

Yang Z, Rannala B. 2014. Unguided species delimitation using DNA sequence data from multiple loci. Mol Biol Evol. 31(12):3125–3135.

Zhang C, Ogilvie HA, Drummond AJ, Stadler T. 2018. Bayesian inference of species networks from multilocus sequence data. Mol Biol Evol. 35(2):504–517.

Zhang W, Dasmahapatra KK, Mallet J, Moreira GRP, Kronforst MR. 2016. Genome-wide introgression among distantly related Heliconius butterfly species. Genome Biol. 17(1):25.

Zhang Y, Teng D, Lu W, Liu M, Zeng H, Cao L, Southcott L, Potdar S, Westerman E, Zhu AJ, et al. 2021. A widely diverged locus involved in locomotor adaptation in Heliconius butterflies. Sci Adv. 7(32):eabh2340.

Zhu T, Flouri T, Yang Z. 2022. A simulation study to examine the impact of recombination on phylogenomic inferences under the multispecies coalescent model. Mol Ecol. 31(10):2814–2829.

