## Supplementary material for "Major patterns in the introgression history of *Heliconius* butterflies": SI figures and texts

Supplementary Information

- Supplementary figures: Figures S1–S20.
- Supplementary tables: Tables S1–S23 (see Excel file).
- Supplementary texts

### Supplementary figures

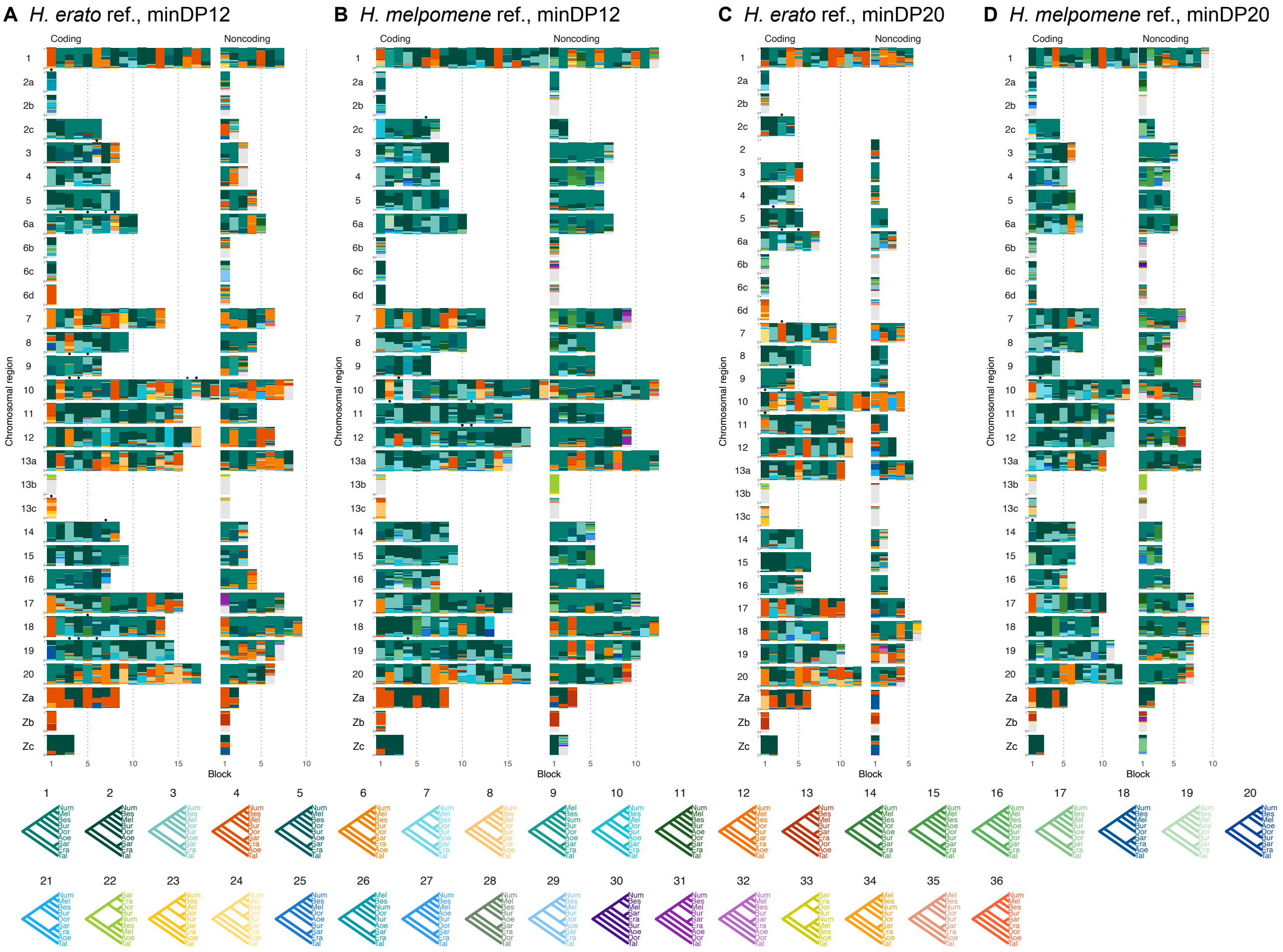

**Figure S1.** Posterior estimates of species trees from four versions of the 'etales-9spp' dataset (see **Methods**). See legend to **Figure 1**.

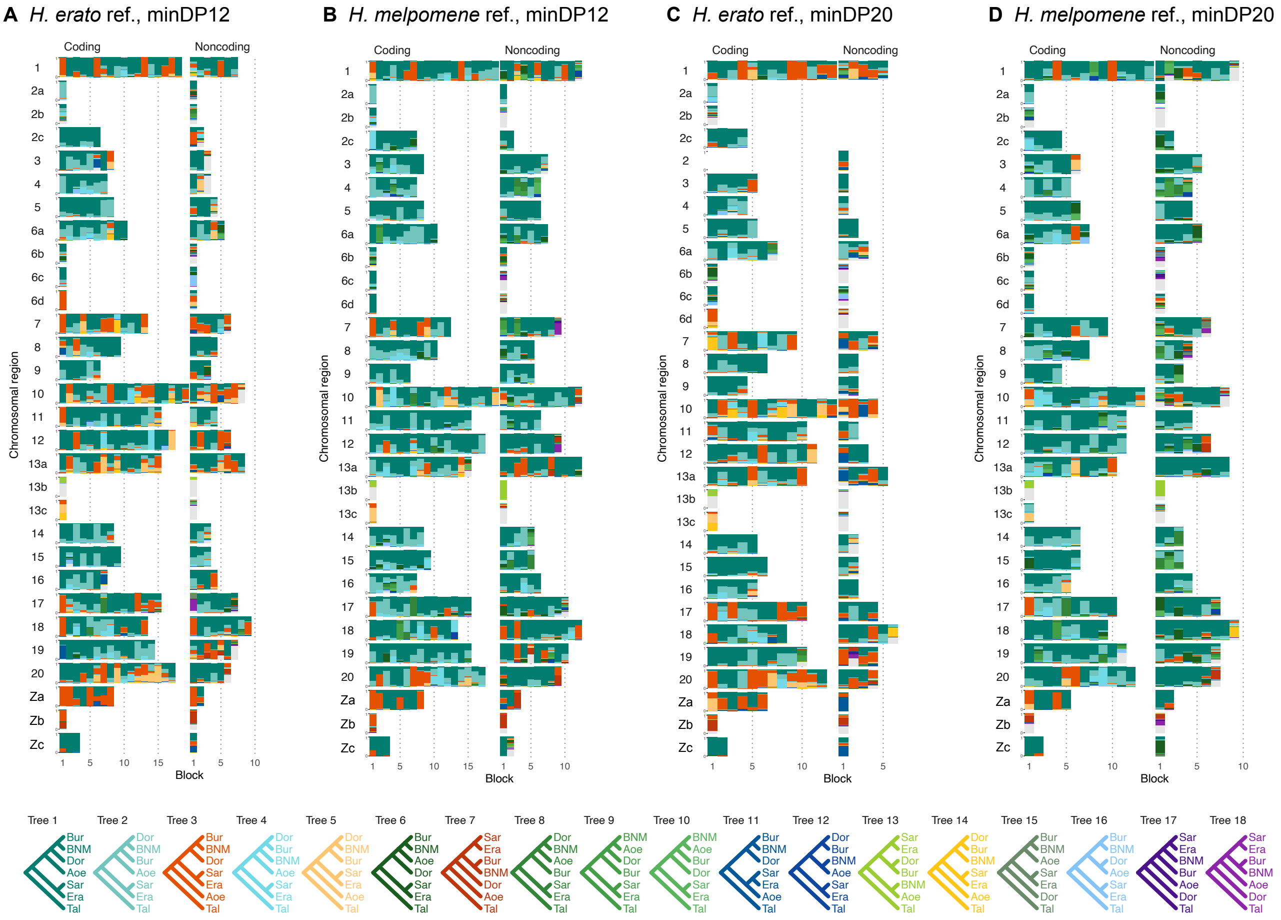

**Figure S2.** Posterior estimates of species trees from four versions of the 'etales-9spp' dataset (see **Methods**) with the clade containing three species (Bes, Num, Mel) in the melpomene-silvaniform clade collapsed as a single species. See legend to **Figure 1**.

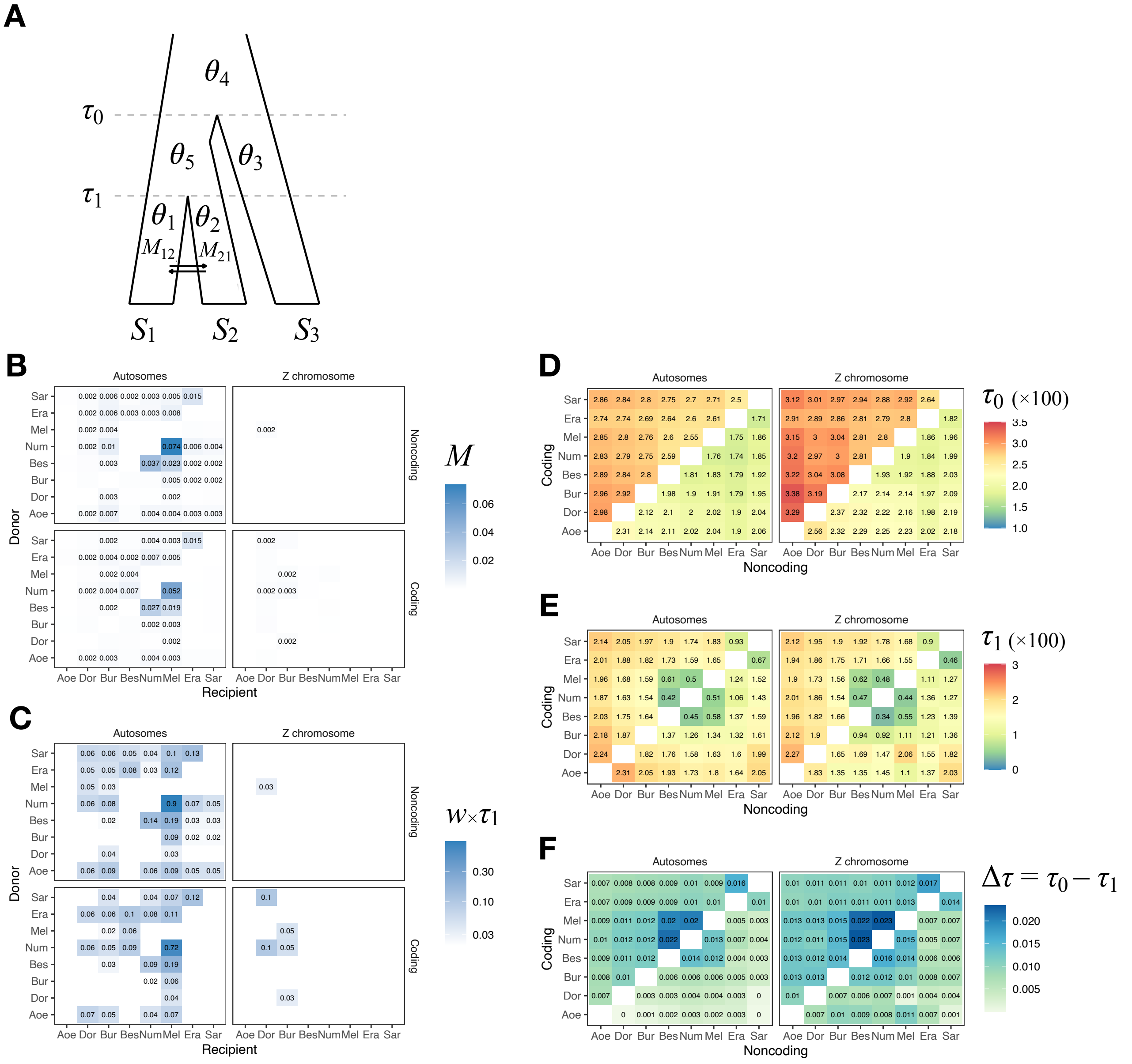


**Figure S3.** Isolation-with-migration (IM) analysis of the 'etales-9spp' dataset using 3S. (A) Model specification in 3S. There are three types of parameters: divergence times ($\tau_1$, $\tau_0$), effective population sizes ($\theta_1$, $\theta_2$, $\theta_4$, and $\theta_5$) and migration rates ($M_{12}$, $M_{21}$). (B) Maximum-likelihood estimates (MLEs) of the pairwise migration rates ($M_{12}$, $M_{21}$), with donor species on the y-axis and recipient species on the x-axis. We used *Eueides tales* as the outgroup ($S_3$ in A). Top and bottom quadrants are results from noncoding and coding loci, respectively. Left quadrants show results from all autosomal loci. Right quadrants show results from all Z chromosome loci (21a+21b+21c). For the number of loci, see **Table S3** (*H. erato* reference, minDP ($d$) = 12). For pairs without displayed numbers, the likelihood ratio test (LRT) was not significant at 0.1%, thereby failing to reject a null model of zero gene flow. (C) Mutation-scaled migration rates ($w_{12} = 4M_{12}/\theta_2$ and $w_{21} = 4M_{21}/\theta_1$) as a measure of expected admixture fraction in the recipient genome. (D) Root age ($\tau_0$). Estimates from coding loci are shown in the upper triangle while estimates from noncoding loci are in the lower triangle. (E) Divergence time of the ingroup species pair ($\tau_1$). (F) Internal branch length ($\Delta\tau = \tau_0 - \tau_1$). Full results are in **Table S8** and **Figure S4**.

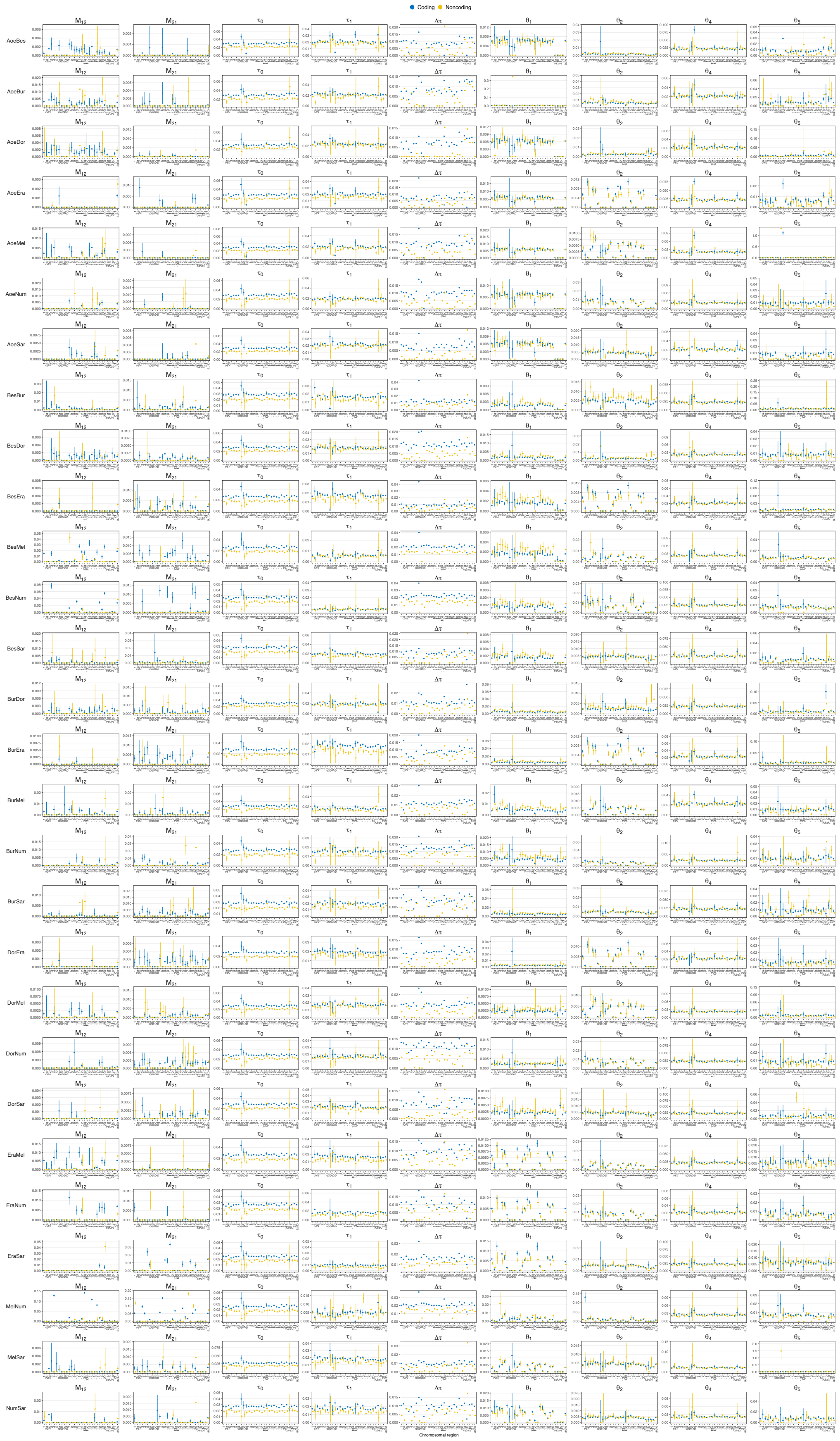


**Figure S4.** MLEs of all parameters in the MSC-M model as well as internal branch lengths ($\Delta\tau = \tau_0 - \tau_1$) by chromosomal region for all pairs of *Heliconius* species in the 'etales-9spp' dataset obtained from 3S analysis. Error bars indicate two standard errors (not shown if standard errors were not reliably estimated). Columns are parameters in the model; rows are species pairs.

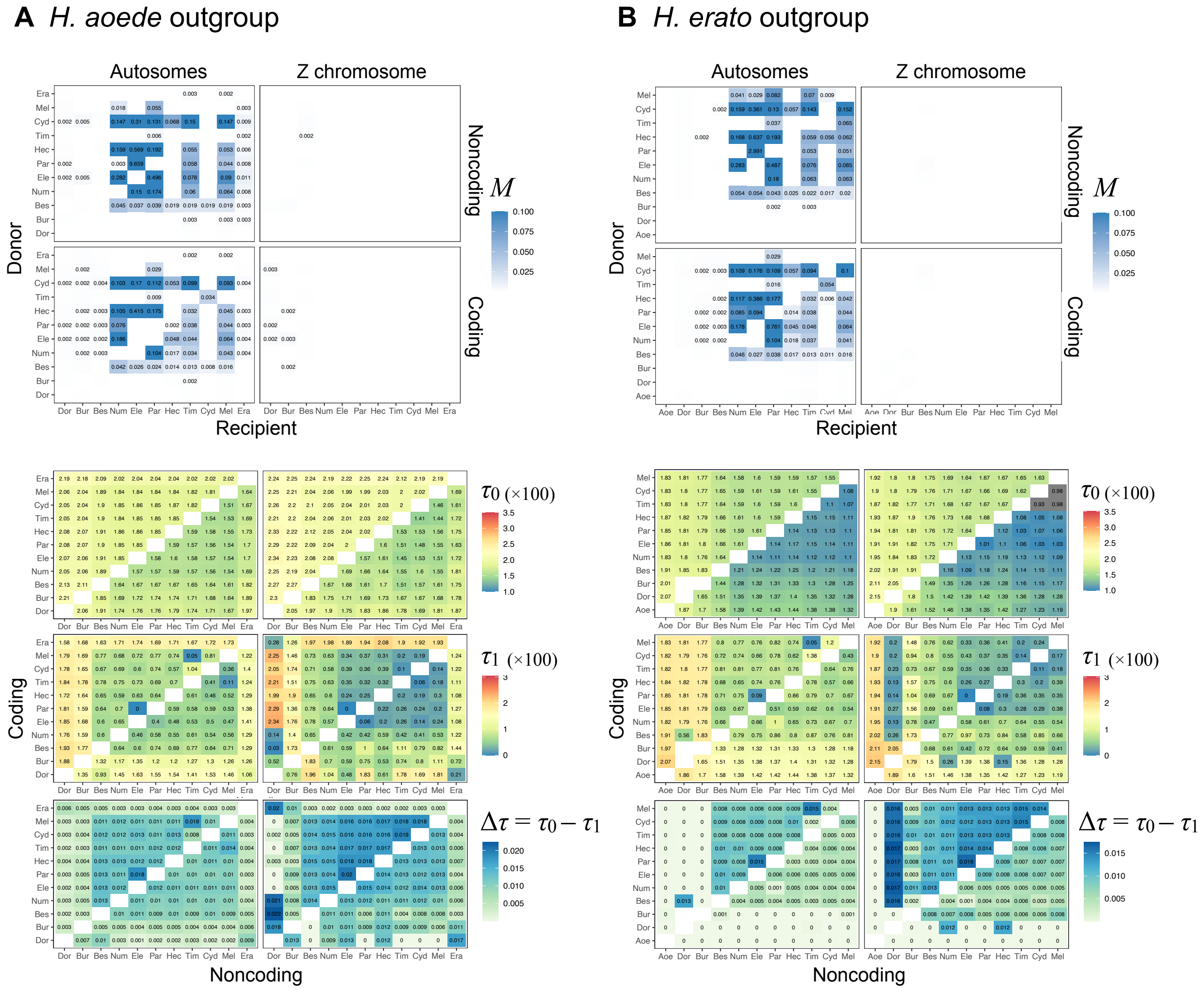


**Figure S5.** MLEs of migration rates ($M$), divergence times ($\tau_0$, $\tau_1$) and the internal branch length ($\Delta\tau = \tau_0 - \tau_1$) from 3s analysis of the 'hmelv25-all' dataset, using (A) *H. aoede* or (B) *H. erato* as an outgroup. See legend to **Figure S3**.

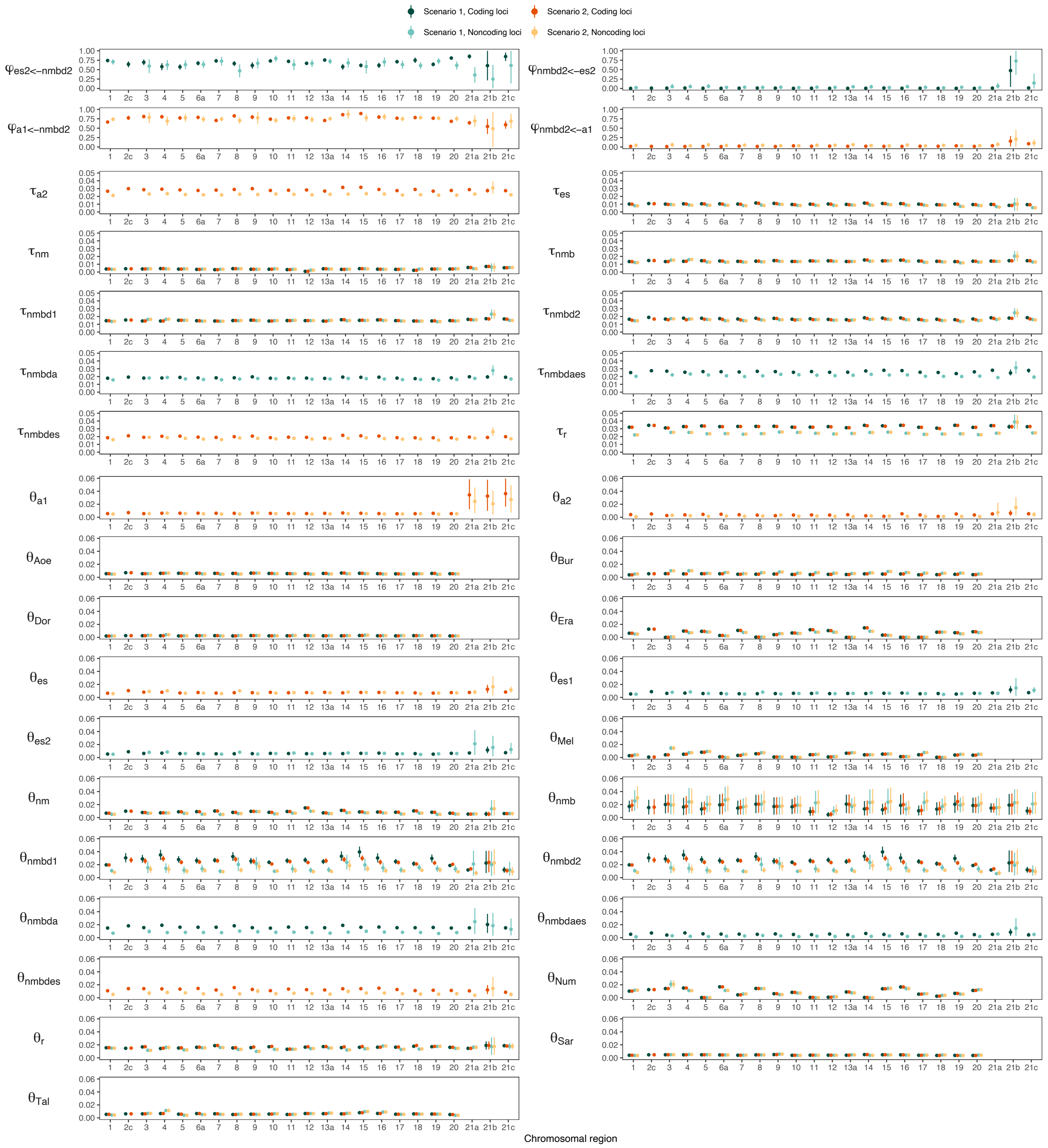


**Figure S6.** Posterior means and 95% HPD intervals of introgression probabilities ($\varphi$), population sizes ($\theta$) and divergence or introgression times ($\tau$) under scenario 1 and scenario 2 MSC-I models in **Figure 1D**,**E** obtained from BPP analysis of the 'etales-8spp' dataset (see **Table S3** for the number of loci). A few chromosomal regions (2a, 2b, 6b, 6c, 6d, 13b, 13c) yielded too few loci (<100) and were excluded.

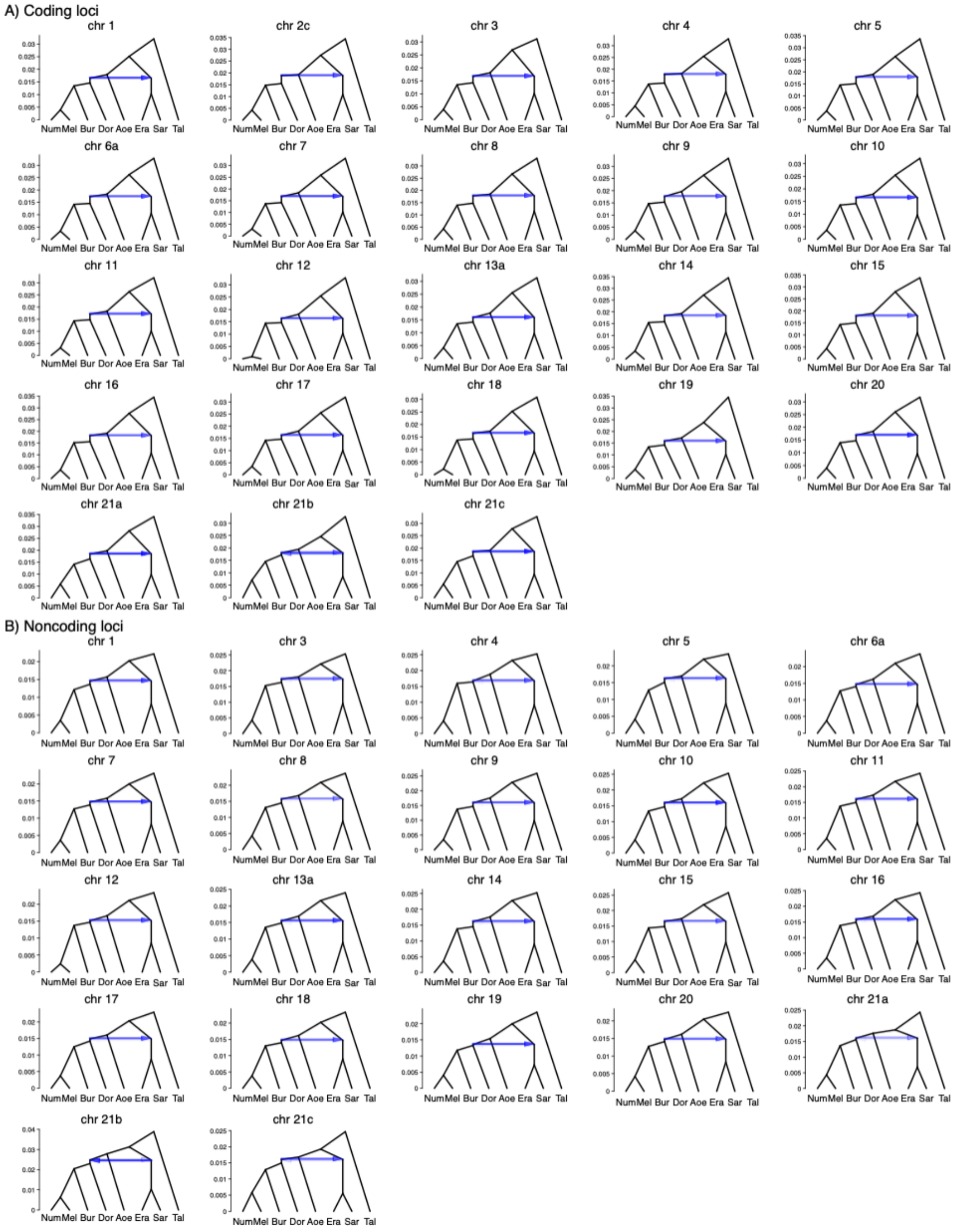


**Figure S7.** Posterior estimates of the species tree under the introgression model in **Figure 1D** (scenario 1: erato-early) obtained from BPP analysis of the 'etales-8spp' dataset for each chromosomal region.

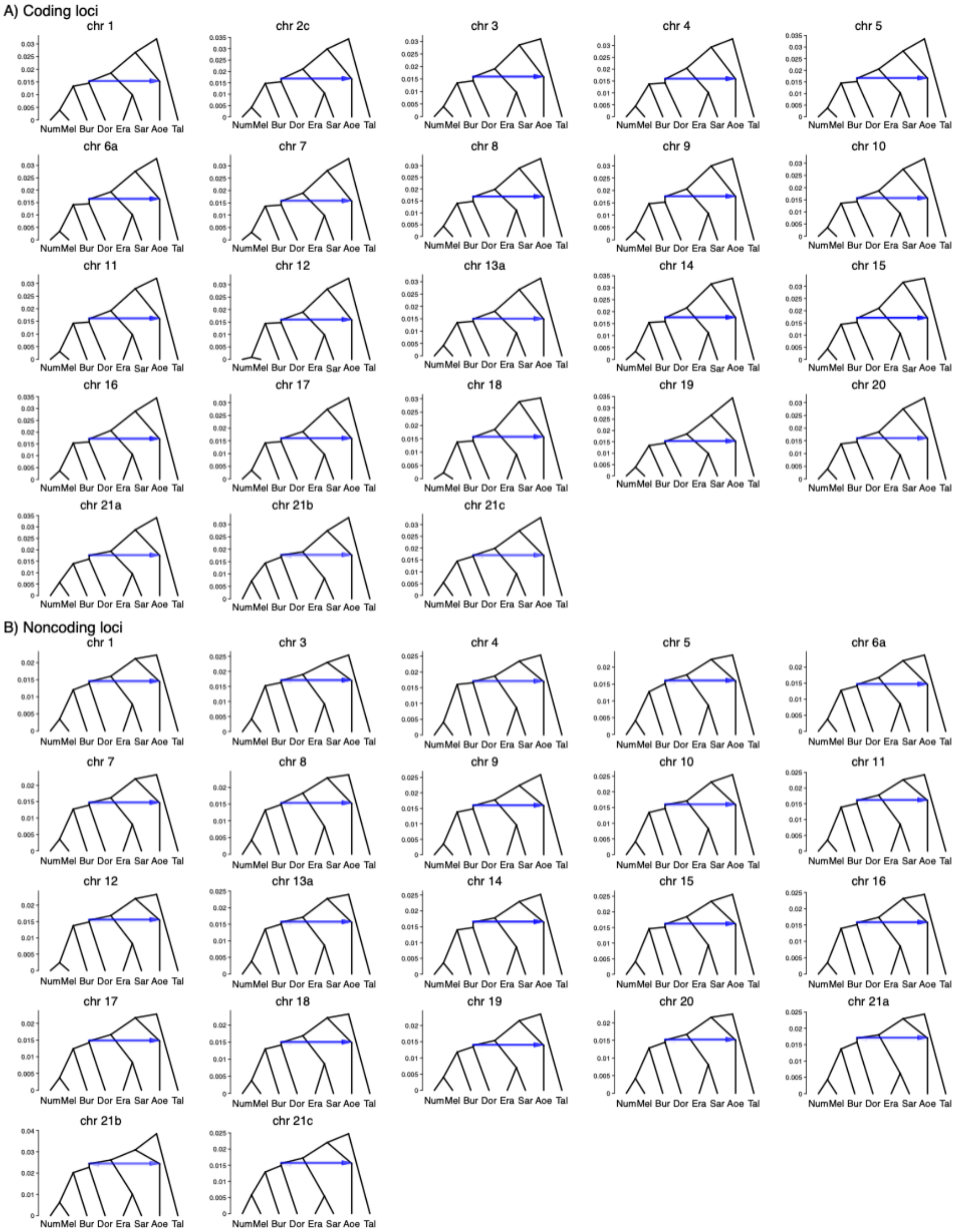


**Figure S8.** Posterior estimates of the species tree under the introgression model in **Figure 1E** (scenario 2: aoede-early) obtained from BPP analysis of the 'etales-8spp' dataset for each chromosomal region.

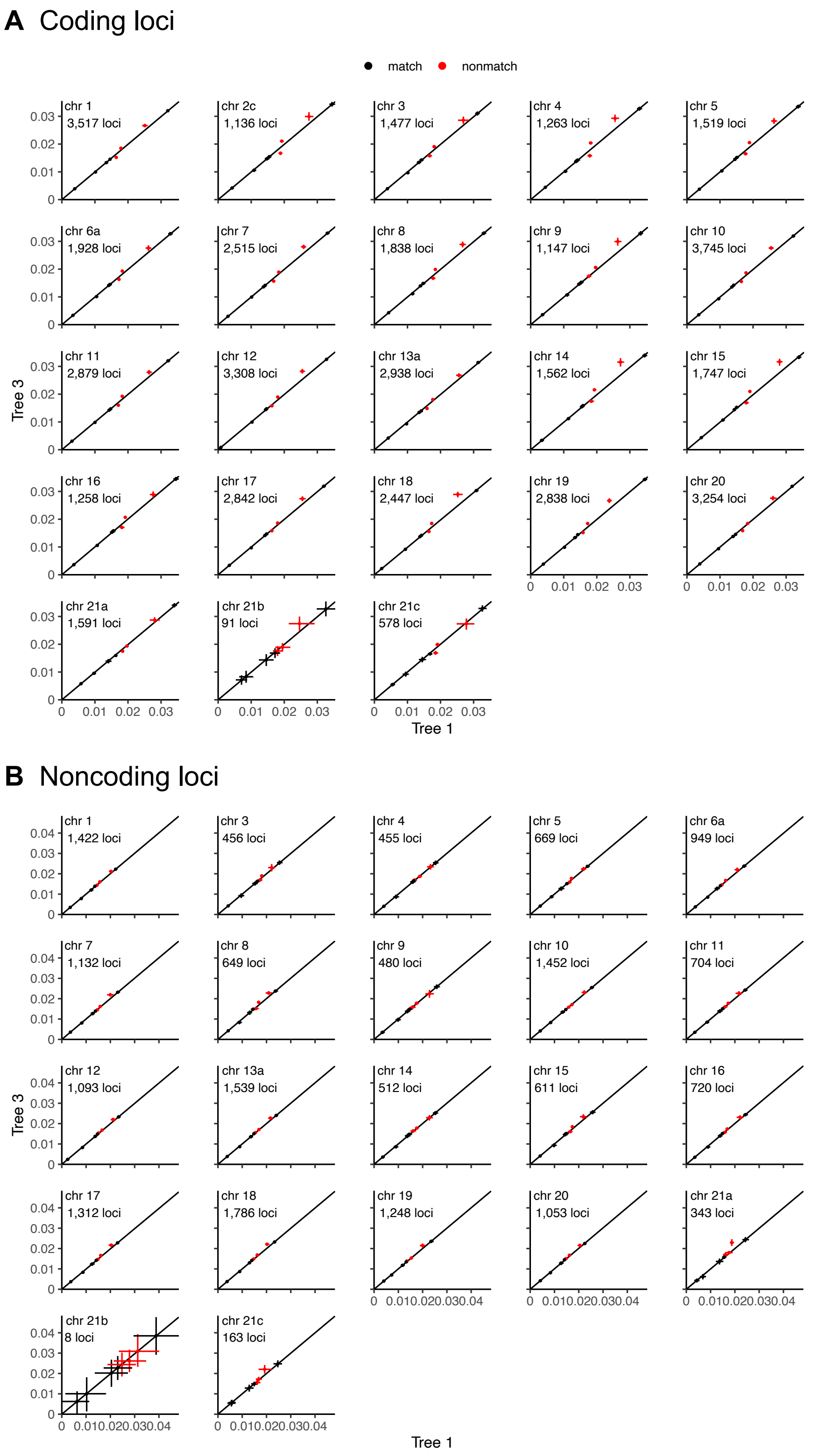


**Figure S9.** Posterior means and 95% HPD intervals of divergence times estimated under the MSC-I model of **Figure 1D** (erato-early, x-axis) versus estimates under the MSC-I model of **Figure 1E** (aoede-early, y-axis) using coding (A) or noncoding loci (B) of the 'etales-8spp' dataset. Red points are divergence times unique to each model.

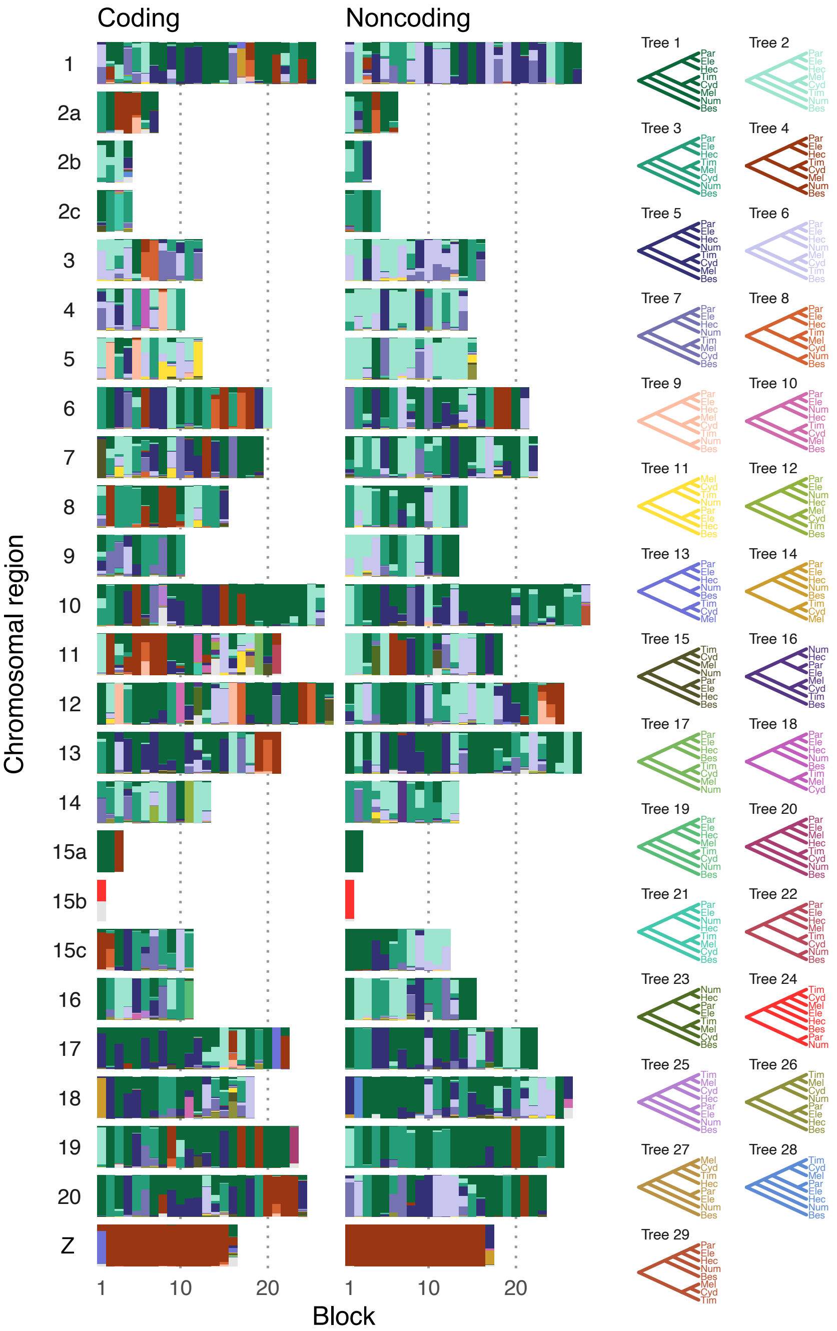


**Figure S10.** Posterior estimates of species trees of the melpomene-silvaniform clade inferred from 200-locus blocks across the genome from the BPP MSC analysis with no gene flow of the 'hmelv25-res' dataset. **Figure 2A** shows a simplified version with fewer trees. See **Tables S11** and **S12**.

**A** *H. aoede* outgroup **B** *H. erato* outgroup

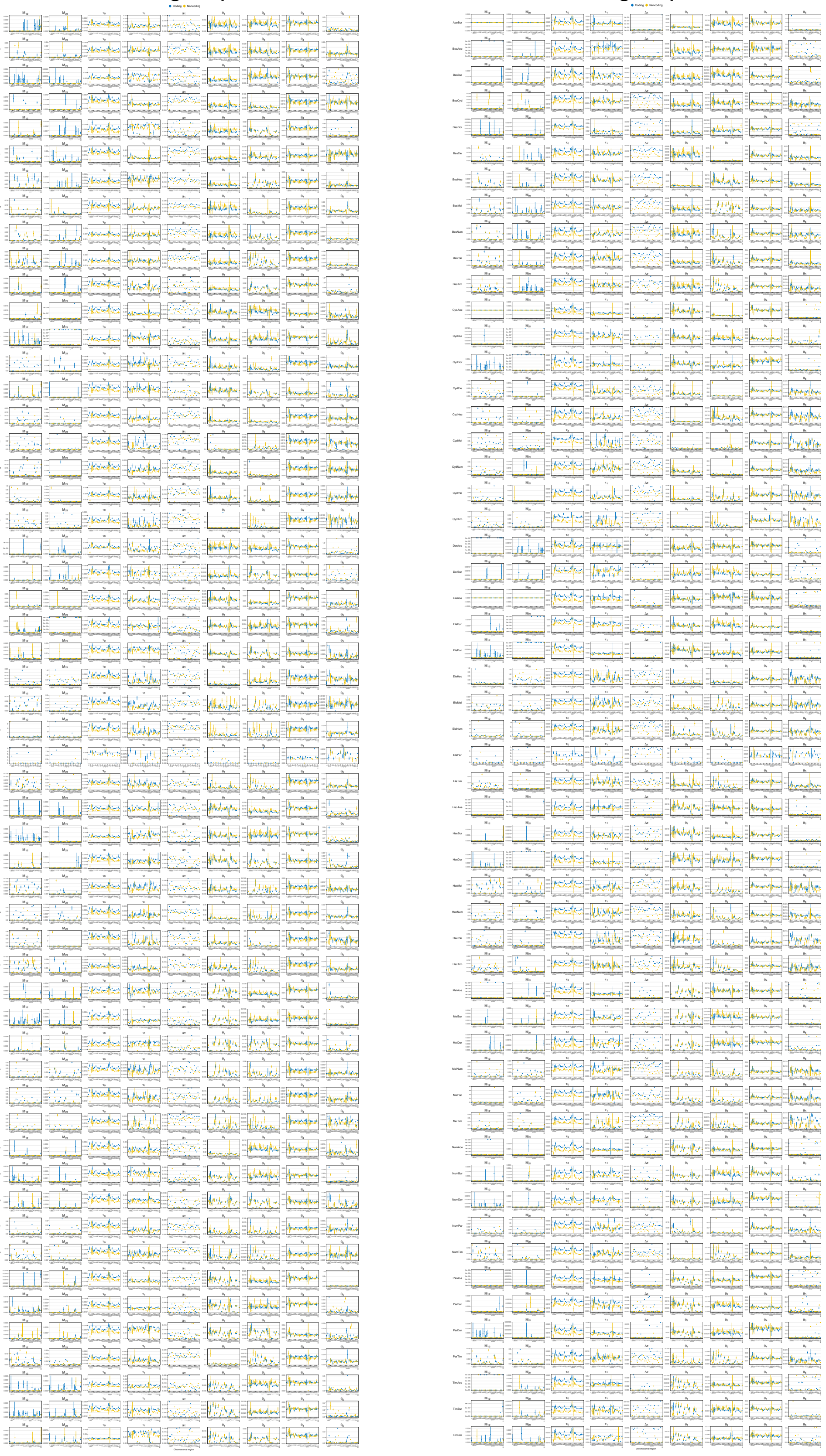

**Figure S11**. Maximum likelihood estimates (MLEs) of all parameters in the MSC-M model as well as internal branch lengths ($\Delta\tau = \tau_0 - \tau_1$) by chromosomal region from species pairs in the 'hmelv25-all' dataset obtained from 3s analysis using (A) *H. aoede* or (B) *H. erato* as an outgroup. Error bars indicate two standard errors (not shown if standard errors were not reliably estimated). Columns are parameters in the model; rows are species pairs. Full results are in **Tables S15** and **S16**.

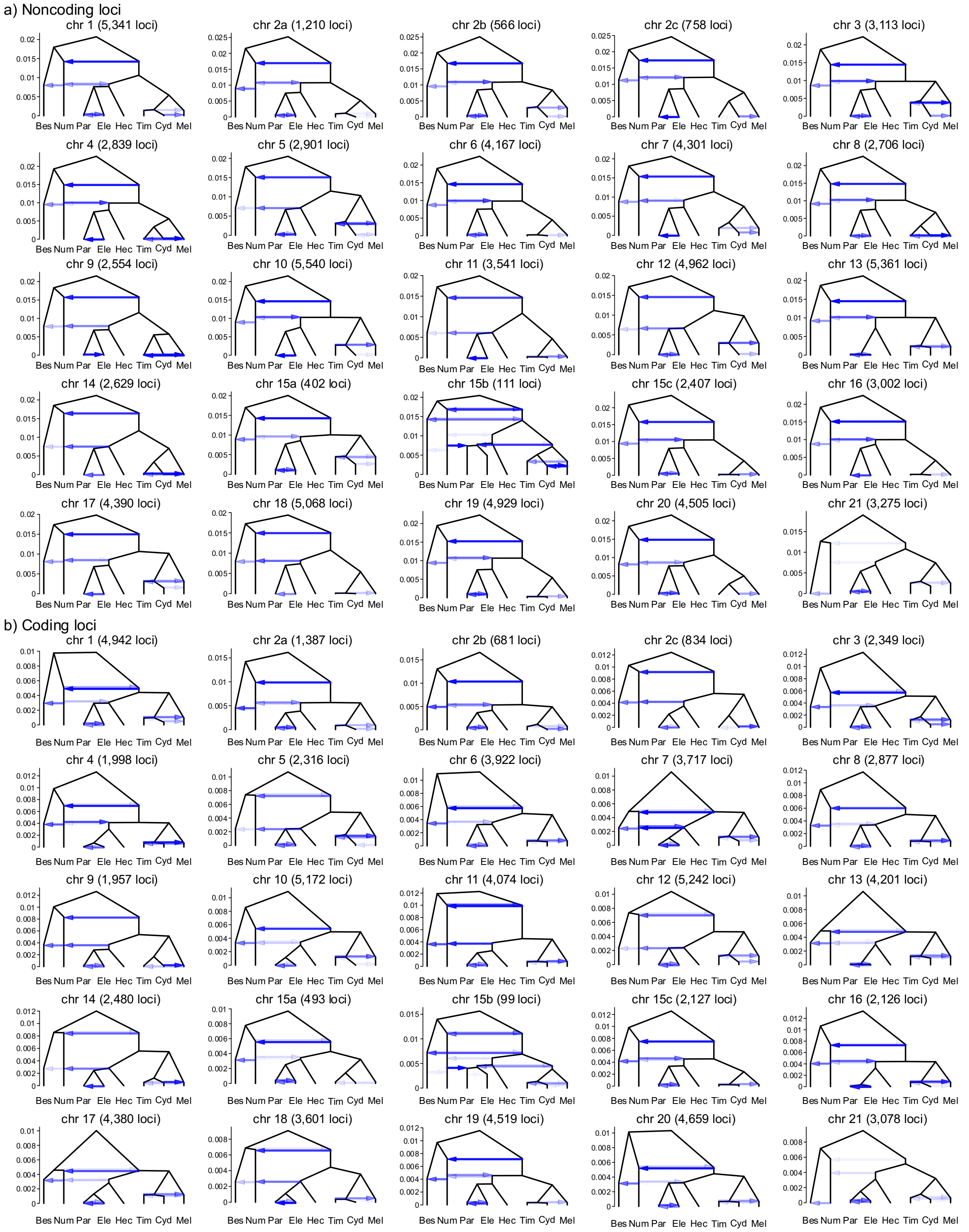


**Figure S12. Estimated introgression history.** Estimated introgression history for each chromosomal region obtained under the model of **Figure 2B** and **Figure S13D** using (A) noncoding and (B) coding loci. Intensity of the horizontal edges is proportional to posterior mean of introgression probability, while the y-axis represents the divergence times in the units of the expected number of mutations per site. For posterior estimates of all parameters, see **Tables S17–S18**.

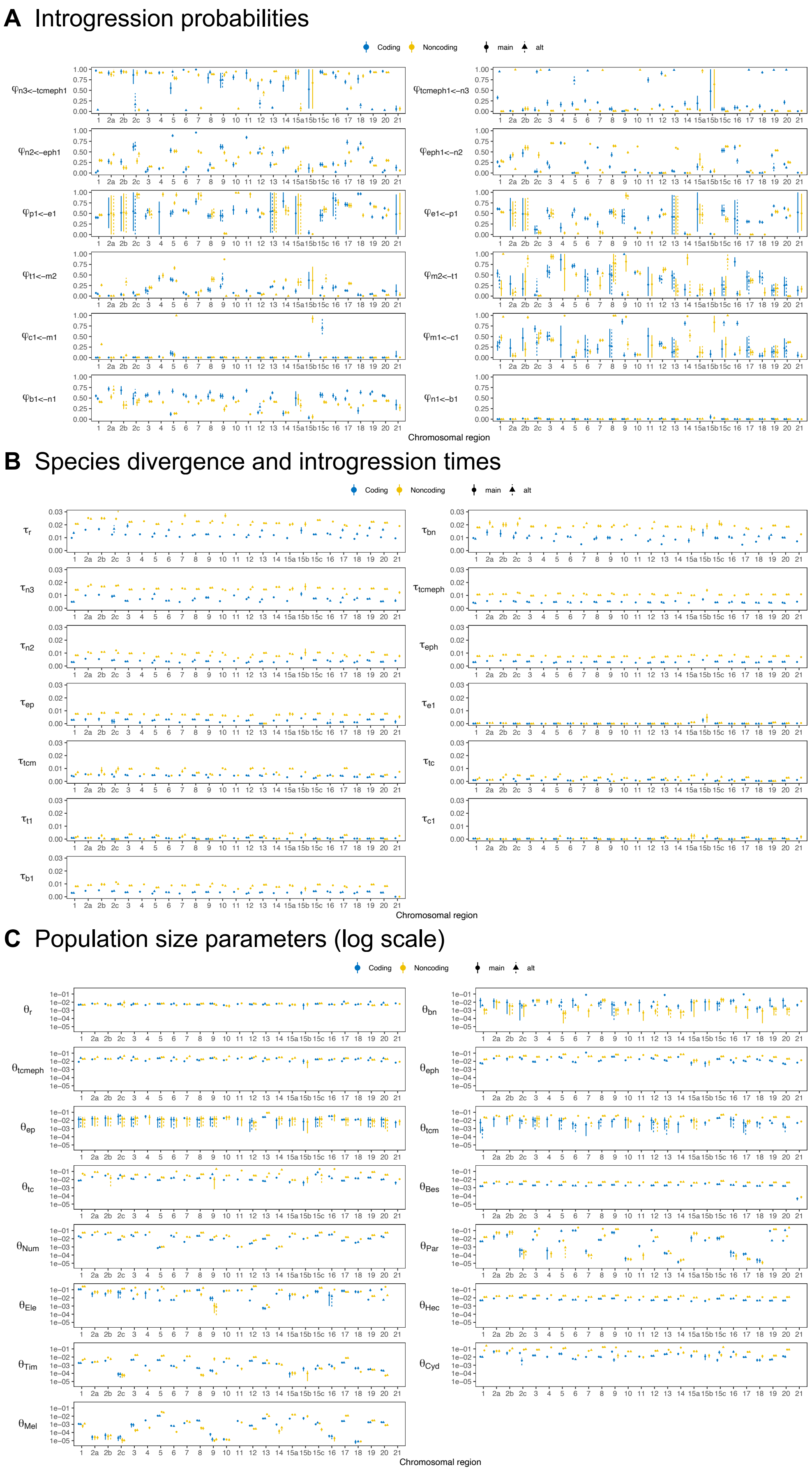
A Introgression probabilities
Coding
Noncoding
main
alt
Chromosomal region
B Species divergence and introgression times
Coding
Noncoding
main
alt
Chromosomal region
C Population size parameters (log scale)
Coding
Noncoding
main
alt
Chromosomal region

**D** Introgression model of the 15b region

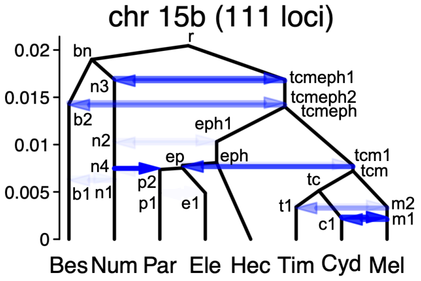


**E** 15b-specific parameters

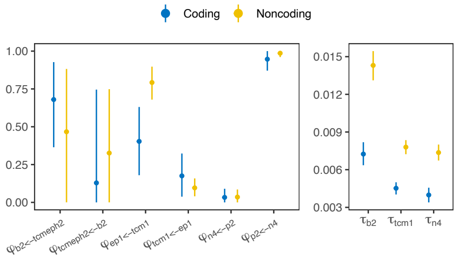


**Figure S13**. Posterior means and 95% HPD intervals of (A) introgression probabilities, (B) divergence or introgression times ($\tau$), (C) population sizes ($\theta$) under the MSC-I model of **Figure 2B**. (D) Introgression model of the chromosome 15b inversion region. (E) Additional introgression time ($\tau$) and probability ($\varphi$) parameters specific to the MSC-I model for the chr 15b inversion region (D).

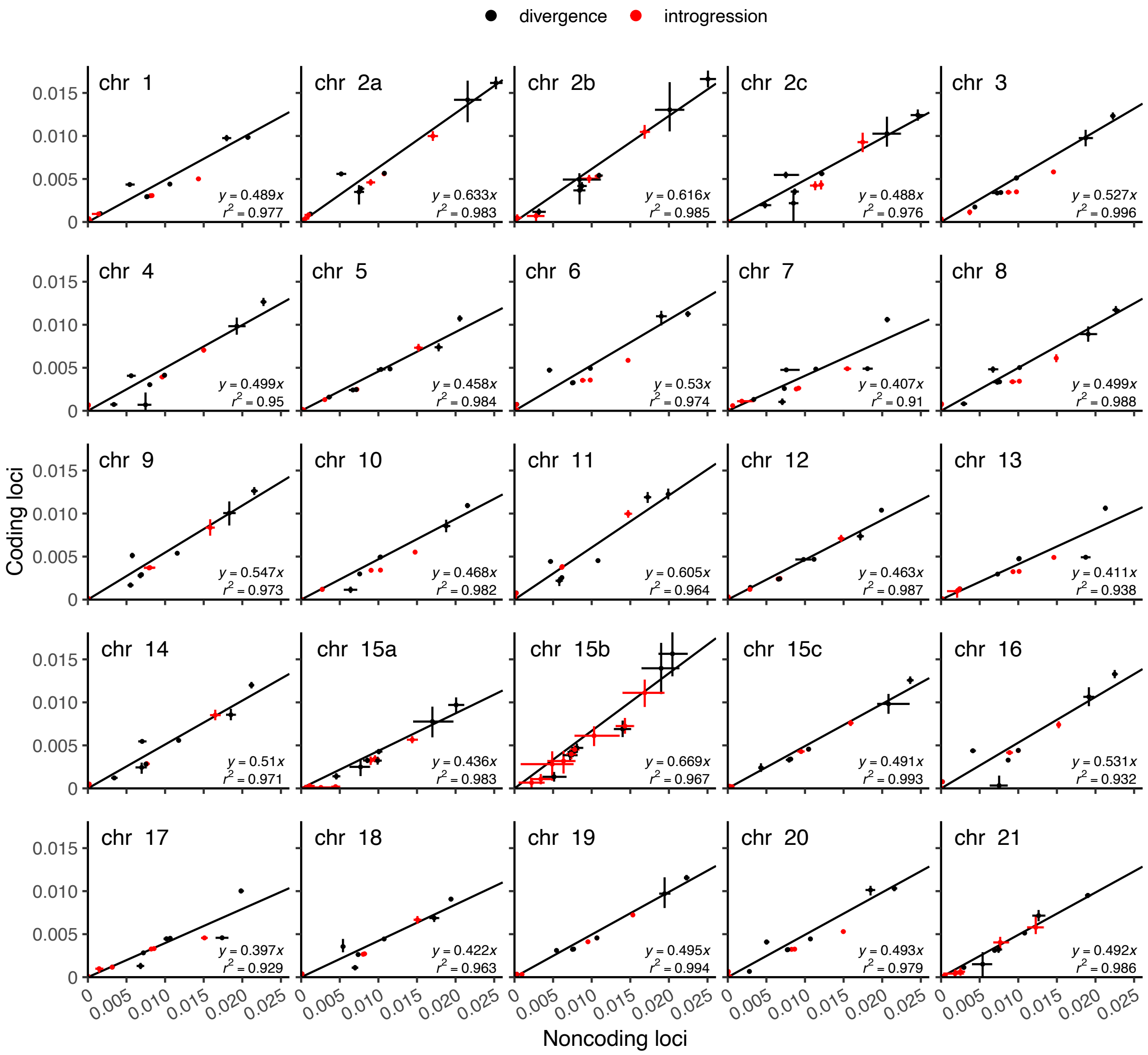


**Figure S14.** Posterior means and 95% HPD intervals of species divergence and introgression times ($\tau$) obtained from coding loci (*y*-axis) versus noncoding loci (*x*-axis) under the MSC-I models of **Figure 2B** (for chromosomal regions other than 15b) and of **Figure S13D** (for 15b). We used the divergence times (black points) only to fit a regression line $y = c\,x$. For the number of loci used, see **Table S12** or **Figure S12**.

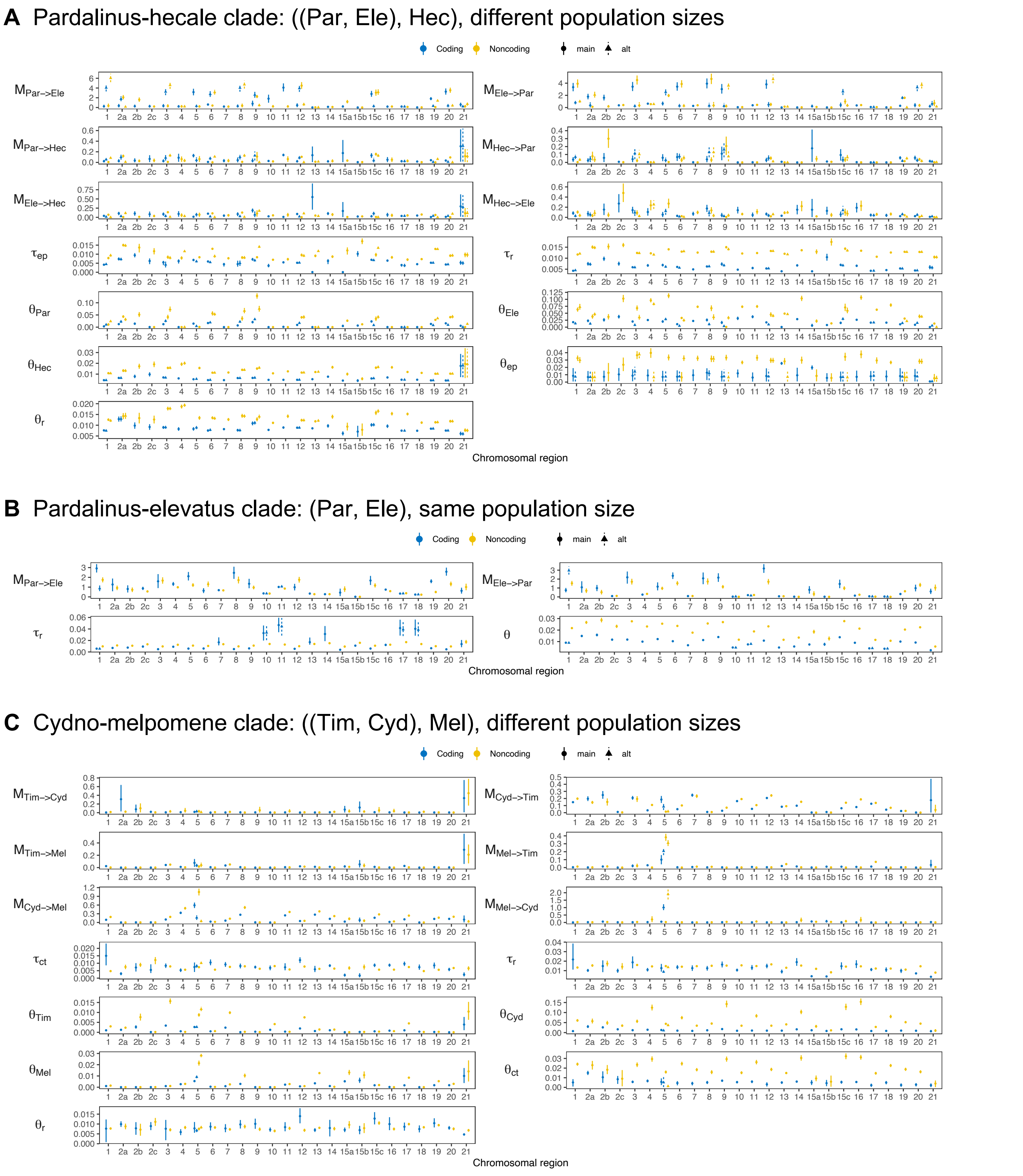


**Figure S15.** Posterior means and 95% HPD intervals of migration rates (*M*), divergence times ($\tau$) and population sizes ($\theta$) under the MSC-M model in of three species obtained using BPP. (A) Pardalinus-hecale clade: ((Par, Ele), Hec). (B) Pardalinus-elevatus clade: (Par, Ele), assuming all three populations have the same size $\theta$. (C) Cydno-melpomene clade: ((Tim, Cyd), Mel). Full results for these three models are given in **Tables S19–S21**.

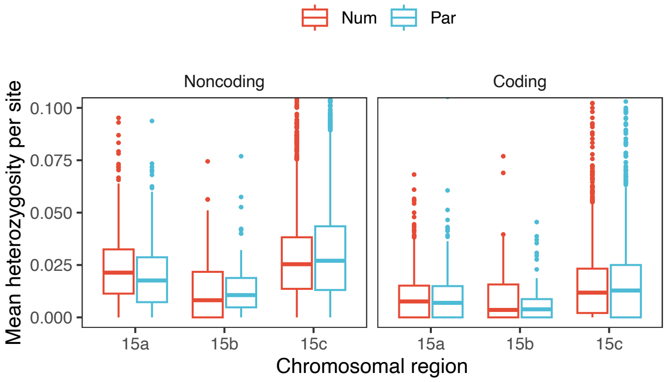


**Figure S16.** Mean heterozygosity per site of the 15b inversion region and flanking regions (15a and 15c) of *H. numata* (Num) and *H. pardalinus* (Par) individuals. Both individuals are homozygotes for the $P_1$ inversion.

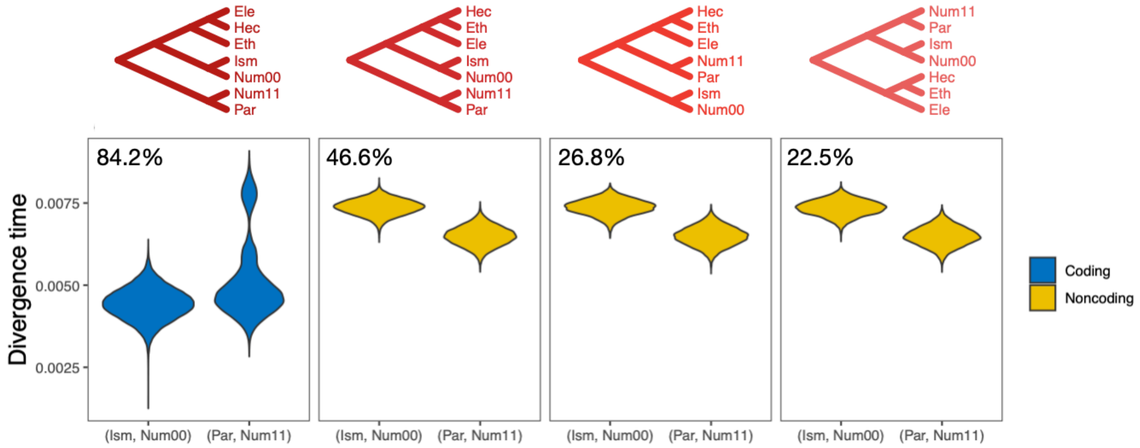


**Figure S17.** Posterior estimates of divergence time for (Ism, Num00) and (Par, Num11) in the 15b region. Only estimates from trees with posterior probabilities >0.1 are shown. Trees are in order of decreasing frequency among coding or noncoding data. See legend to **Figure 3** for species code.

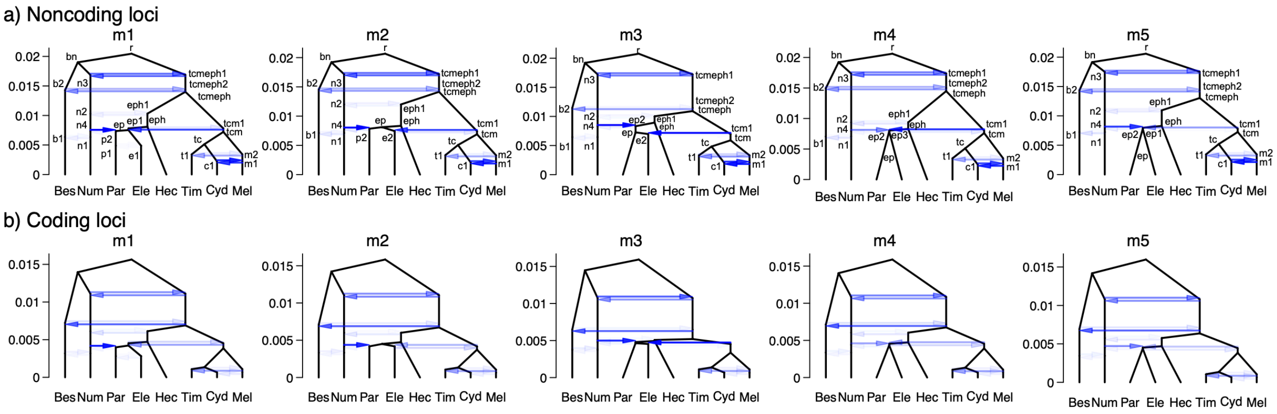


**Figure S18.** Estimated introgression history under five introgression models of the 15b region. See **Table S23** for posterior estimates of all parameters under each model.

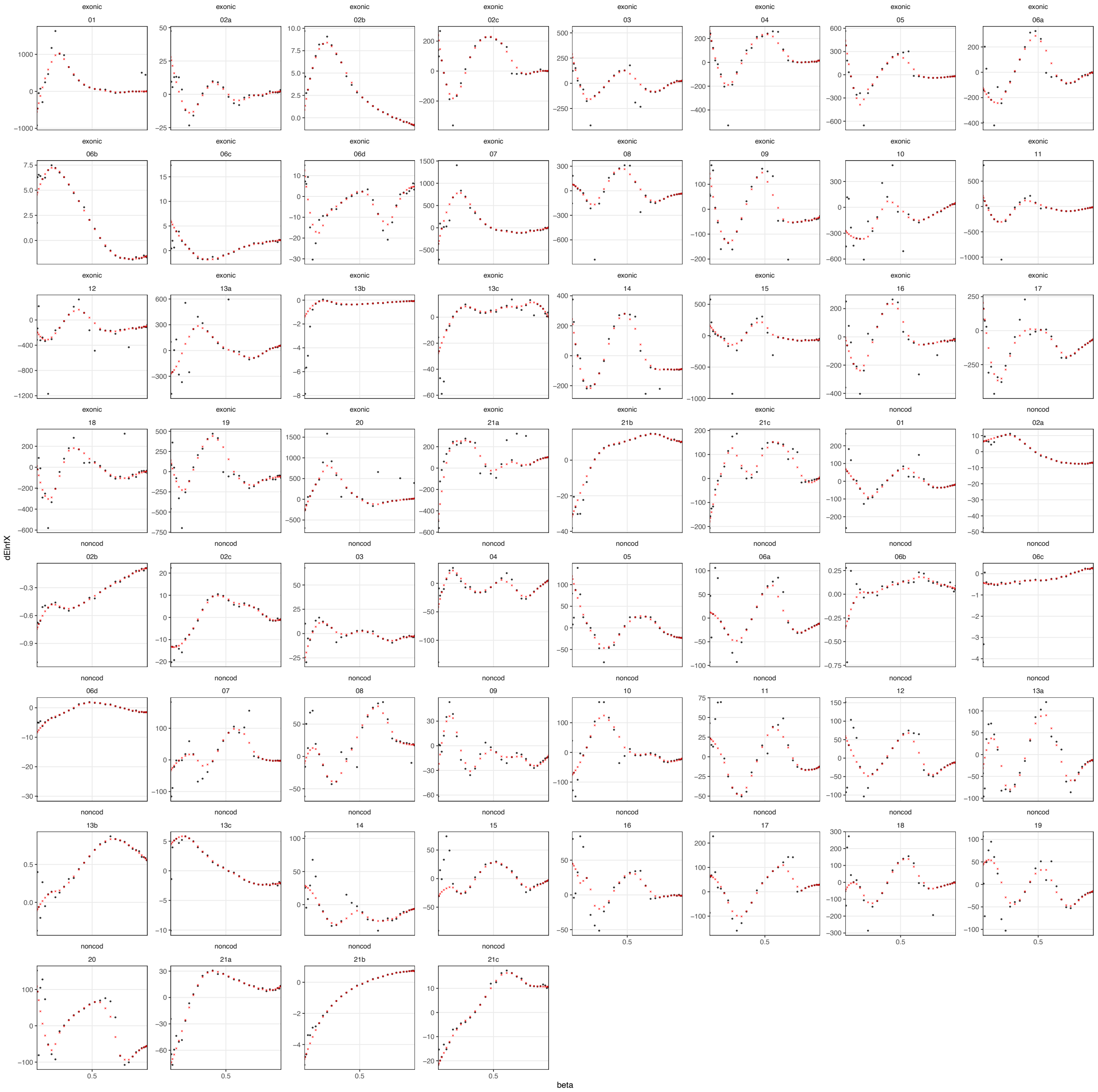

**Figure S19.** Raw (black) and adjusted (red) estimates of the difference in the mean log likelihood from the two models (y-axis). Raw and adjusted estimates of the Bayes factor are in **Table S11**.

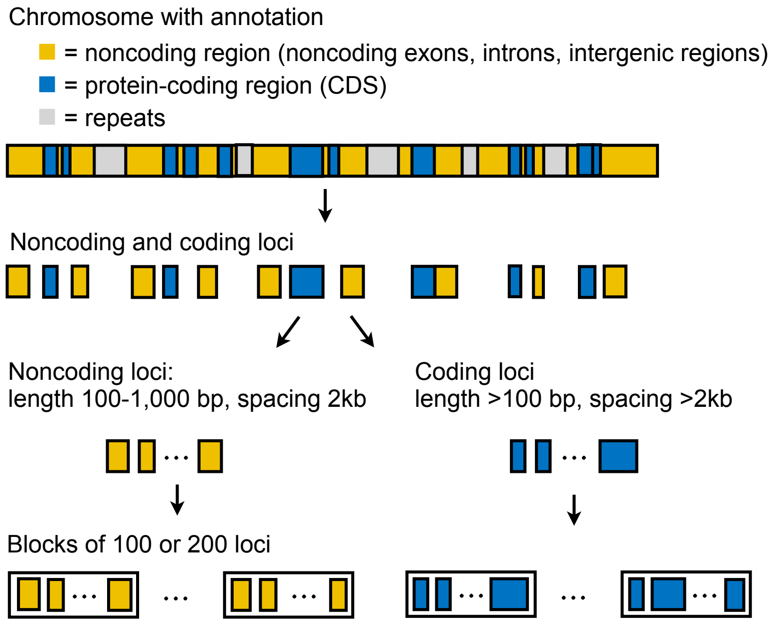


**Figure S20.** Generation of multilocus data. Coordinates of coding and noncoding loci were obtained from a reference genome with an annotation of coding sequences (CDS).

### Supplementary tables

**Table S1.** Sample information.

**Table S2.** Coordinates of chromosome inversion regions considered in this study.

**Table S3.** Number of coding and noncoding loci by chromosomal region in four versions of the 'etales-9spp' dataset.

**Table S4.** Proportions of posterior species tree estimates (MAP trees) from BPP MSC analysis without gene flow of all four versions of the 'etales-9spp' dataset in blocks of 200 loci, with BNM clade (*H. besckei*, *H. numata*, *H. melpomene*) collapsed into a single tip, summarized into major regions: auto (all autosomes excluding inversion regions), the Z chromosome (chr 21, excluding the inversion region), and individual inversion regions (2b, 6b, 6c, 13b and 21b). $n$ = number of blocks. Tree indices correspond to those in **Figure S2**. A full list of MAP trees for each chromosomal region is **Table S5**. Full results without BNM lumping are in **Tables S6–S7**.

**Table S5.** Proportions of MAP trees from BPP MSC analysis without gene flow of all four versions of the 'etales-9spp' dataset in blocks of 200 loci, with BNM clade collapsed into a single tip, summarized by chromosomal region. Genome-wide summaries are given in **Table S4**.

**Table S6.** Full results from BPP MSC analysis without gene flow of the 'etales-9spp' dataset, summarized into major regions of the genome. See also legend to **Table S4**. Trees are in a decreasing order of the total frequencies across all four versions of the dataset. Tree indices correspond to those in **Figure S1**.

**Table S7.** Full results from BPP MSC analysis without gene flow of the 'etales-9spp' dataset, summarized by chromosomal region. See legend to **Table S6**.

**Table S8.** Maximum likelihood estimates (MLEs) of parameters under the isolation-with-migration (IM) model obtained from 3S for each of the 28 pairs of species in the 'etales-9spp' dataset (*H. erato* reference, minDP12), using *Eueides tales* (Tal) as an outgroup. We used the likelihood ratio test (LRT) statistic to test if the model with gene flow (M2) was preferred to the model without gene flow (M0), using the *p*-value threshold of 1%. The 'auto' dataset used all autosomal loci excluding the inversion regions (see **Table S3** for the number of loci). Plots of migration rate and divergence time estimates are in **Figure S3**. Plots of all parameter estimates with confidence intervals are in **Figure S4**.

**Table S9.** Posterior means and 95% HPD intervals (in parentheses) of parameters from the BPP MSC-I model in **Figure 1D** (scenario 1) using noncoding and coding loci for each chromosomal region of 'etales-8spp' dataset (see **Table S1**). Multiple posterior peaks, if present, are reported, e.g. there are two peaks, denoted chr1 and chr1-alt, from the coding loci in chromosome 1. Plots of these estimates with 95% HPD intervals are in **Figure S6**. Plots of species tree with estimated divergence times, introgression times and introgression probabilities are in **Figure S7**.

**Table S10.** Posterior means and 95% HPD intervals (in parentheses) of parameters from the BPP MSC-I model in **Figure 1E** (scenario 2) using noncoding and coding loci for each chromosomal region. Multiple posterior peaks, if present, are reported, e.g. there are two peaks, denoted chr1 and chr1-alt, from the coding loci in chromosome 1. Plots of these estimates with 95% HPD intervals are in **Figure S6**. Plots of species tree with estimated divergence times, introgression times and introgression probabilities are in **Figure S8**.

**Table S11.** Bayes factor $B = M_{1E}/M_{1D}$ for comparing the two MSC-I models in **Figure 1D** ($M_{1E}$; scenario 1) and **Figure 1E** ($M_{1D}$; scenario 2), calculated for coding and noncoding loci for each chromosomal region using thermodynamic integration with 32 Gaussian quadrature points. $M_{1E}$ (scenario 2) is preferred if $B > 100$, $M_{1D}$ (scenario 1) is preferred if $B < 0.01$, and the test is not significant (n.s.) otherwise (i.e. $100 < B < 0.01$). The last three columns are adjusted values of

Bayes factor to account for numerical instability of the thermodynamic integration estimates (see Methods and **Figure S19**).

**Table S12.** Number of coding and noncoding loci in each chromosomal region in the hmelv25-res' dataset used in BPP analysis and the hmelv25-all' dataset use in 3S analysis.

**Table S13.** Proportions of MAP trees, with minimum, median and maximum posterior probabilities shown in parentheses, from BPP MSC analysis without gene flow of the 'hmelv25-res' dataset in blocks of 200 loci, summarized into four major regions: auto (all autosomes excluding 2b and 15b), 2b and 15b inversion regions, and chromosome 21 (Z chromosome). Trees are in a decreasing order of combined frequency across all chromosomes. Tree indices correspond to those in **Figure 10**.

**Table S14.** Proportions of MAP trees, with minimum, median and maximum posterior probabilities shown in parentheses, from BPP MSC analysis without gene flow in blocks of 200 loci, summarized by chromosomal region. See legend to **Table S13**.

**Table S15.** MLEs of parameters under the MSC-M model obtained from 3S for each of the 55 pairs of species in the 'hmelv25-all' dataset using *H. aoede* as an outgroup. We used the LRT statistic to test if the model with gene flow (M2) was preferred to the model without gene flow (M0), using the *p*-value threshold of 1%. See **Figure S11A** for plots of these estimates. The 'auto' setting used all autosomal loci excluding the inversion regions. See **Table S12** for the number of loci.

**Table S16.** MLEs of parameters under the MSC-M model obtained from 3S for each of the 55 pairs of species in the 'hmelv25-all' dataset using *H. erato* as an outgroup. See legend to **Table S15**. See **Figure S11B** for plots of these estimates.

**Table S17.** Posterior means and 95% HPD intervals (in parentheses) of parameters obtained from the BPP MSC-I model of **Figure 2B** using noncoding loci from each non-15b chromosomal region, and under the MSC-I model of **Figure 13D** for the chromosome 15b inversion region. Plots of the estimates are in **Figure 13**. 'n/a' means the parameter does not exist in the model.

**Table S18.** Posterior means and 95% HPD intervals (in parentheses) of parameters from BPP MSC-I models in **Figure 2B** and **Figure 13D** using coding loci. See legend to **Table S17**.

**Table S19.** Posterior means and 95% HPD intervals (in parentheses) of parameters from BPP MSC-M models for the pardalinus-hecale clade: ((Par, Ele), Hec), with six continuous migration rates ($M = Nm$) among the three species, two species divergence times ($\tau$) and five population sizes ($\theta$). Plot of the estimates are in **Figure S15A**.

**Table S20.** Posterior means and 95% HPD intervals (in parentheses) of parameters obtained from BPP MSC-M models for the pardalinus-elevatus clade: (Par, Ele), with two continuous migration rates ($M = Nm$). All populations were assumed to have the same population size ($\theta$). Plot of the estimates are in **Figure S15B**.

**Table S21.** Posterior means and 95% HPD intervals (in parentheses) of parameters from BPP MSC-M models for the cydno-melpomene clade: ((Tim, Cyd), Mel), with six continuous migration rates ($M = Nm$) among the three species, two species divergence times ($\tau$) and five population sizes ($\theta$). Plot of the estimates are in **Figure S15C**.

**Table S22.** Proportions of MAP trees, with minimum, median and maximum posterior probabilities shown in parentheses, from BPP MSC analysis without gene flow of the 'silv-chr15' dataset in blocks of 200 loci or 100 loci, summarized by three regions: the inversion region (15b) and flanking regions (15a and 15c). Trees are in decreasing order of combined frequency across all chromosomal regions and from both options of block size.

**Table S23.** Posterior means and 95% HPD intervals (in parentheses) of parameters obtained from five BPP MSC-I models in **Figure S18**.

### Supplementary text

#### Major introgression patterns in the melpomene-silvaniform clade inferred using 3s and BPP

To assess the possibility of introgression that might support the autosome-majority tree or the Z chromosome tree, we estimate pairwise gene flow rates under the MSC-M model between each pair of species in the melpomene clade as well as *H. burneyi*, *H. doris* and *H. erato* using the program 3s (Zhu and Yang 2012; Dalquen et al. 2017). This implementation allows for continuous bidirectional gene flow between two species since divergence. We performed maximum likelihood estimation and likelihood ratio tests (LRT) to determine if a allowing for bidirectional gene flow (model M2) is a better fit than without gene flow (model M0). An outgroup species was used to improve the power of the test but it was not involved in gene flow. We used *H. aoede* as an outgroup in all species pairs. We find gene flow to be prevalent in the autosomes but almost absent in the Z chromosome (**Figure S5A**). In particular, we find no strong evidence of gene flow in the Z chromosome between *H. numata* and *H. besckei*, which is required to explain the Z chromosome tree if the autosome-majority tree is the true species tree. Taken together, these results suggest that the Z chromosome tree represents the true species branching order, and autosomal trees are a result of extensive introgression throughout the history of this group. Using *H. erato* as an outgroup instead of *H. aoede* leads to the same conclusion (**Figure S5B**), although the internal branch length becomes zero whenever one of the ingroup species is *H. aoede*. A full list of estimates from all species pairs is in **Figure S11** and **Tables S15** and **S16**.

Next, we formulated two models to capture major introgression events in the group, using the Z chromosome tree as a base tree (**Figure 2B,C**). The first model is the main model used for most parts of the genome and had six pairs of bidirectional introgression (**Figure 2B**): two for explaining different positions of *H. numata* (nodes n3–tcmeph1 and n2–eph1, relating trees 1–3, tree 4 and trees 5–7), one for *H. pardalinus*–*H. elevatus* gene flow, which is on-going in sympatric populations (p1–e1) (Rosser et al. 2019), two for gene flow involving *H. melpomene* with *H. cydno* (m1–c1) and with *H. timareta* (m2–t1) (explaining variation among trees 1–3 and trees 5–7), and one for the possibility of *H. besckei* and *H. numata* (b1–n1) as suggested by the relationship between the Z chromosome tree and autosome trees. To keep the model tractable, we excluded potential *H. cydno*–*H. timareta* introgression in the model because these species are presently allopatric and signals of gene flow between them are likely to be via *H. melpomene*. We also assumed that population size did not change after introgression. This model has 12 introgression probabilities ($\varphi$), 13 species divergence times and introgression times ($\tau$) and 15 population size parameters ($\theta$), a total of 40 parameters. The second model is for the chromosome 15 inversion region (15b), which had three additional pairs of introgression edges, with 9 additional parameters (**Figure 13D**). We estimated those parameters for coding and noncoding loci of each chromosomal region using BPP (Flouri et al. 2020).

Our estimated introgression histories are largely consistent across the genome and between coding and noncoding regions (**Figures S13** and **S14** and **Tables S17** and **S18**). This pattern agrees with blockwise variation in species relationships (estimated without gene flow, **Figure 2A**). Gene flow was almost absent in the Z chromosome and consistent with our estimates under the MSC-M model.

- The two introgression events involving *H. numata* that resolve majority autosomal relationships (trees 1–3, 5–7) and the Z chromosome tree (tree 4) were estimated to have high probabilities except for on the Z chromosome and in the chromosome 15b inversion region (**Figure S13A**). First, for the oldest introgression between *H. numata* and the common ancestor of the cydno-melpomene + pardalinus-hecale clade, the introgression is estimated to be unidirectional into *H. numata*, with probability ($\varphi_{\text{n3}\leftarrow\text{tcmeph1}}$) ~0.85 in most parts of the genome while the introgression in the opposite direction was almost absent ($\varphi_{\text{tcmeph1}\leftarrow\text{n3}} < 0.05$). Second, the subsequent introgression between *H. numata* and the common ancestor of the pardalinus-hecale clade is estimated to be bidirectional in many chromosomes, with probabilities ($\varphi_{\text{n2}\leftarrow\text{eph1}}$ and $\varphi_{\text{eph1}\leftarrow\text{n2}}$) ~0.3 on average in both directions, although values of these probabilities varied considerably among chromosomes (**Figure S13A**).

- Within the pardalinus-hecale clade, we estimate introgression probabilities between the sister species *H. pardalinus* and *H. elevatus* ($\varphi_{e1\leftarrow p1}$ and $\varphi_{p1\leftarrow e1}$) to be ~0.5 in both directions in most chromosomes, although there is some variation across the genome. The time of this introgression ($\tau_{e1}$ or $\tau_{p1}$) is essentially zero (**Figure S13A** and **Tables S17** and **18**), consistent with the fact that these two species still hybridize today (Mallet et al. 2007). In some chromosomes such as 13 and 21, posterior estimates of ~0.5 had wide intervals (**Figure S13A**). The uncertainty may be due to weak evidence for gene flow, even with non-zero posterior means.
- Within the cydno-melpomene clade, both *H. melpomene*–*H. cydno* and *H. melpomene*–*H. timareta* introgression is estimated to be highly asymmetrical, with *H. melpomene* being the main recipient in both cases, and with the introgression probabilities varying widely across the genome (**Figure S13A**). The *H. cydno* → *H. melpomene* introgression is estimated to be strictly unidirectional, with introgression probability ($\varphi_{m1\leftarrow c1}$) of ~0.35 in noncoding regions, ~0.25 in coding regions, and zero in the opposite direction ($\varphi_{c1\leftarrow m1}$). For *H. melpomene*–*H. timareta*, the introgression probability for *H. timareta* → *H. melpomene* ($\varphi_{m2\leftarrow t1}$) is highly variable across the genome, with values ranging from close to 0 to almost 1; the opposite direction ($\varphi_{t1\leftarrow m2}$) had low probabilities (<0.1) for most chromosomes.
- Surprisingly, we found strictly unidirectional introgression from *H. numata* into *H. besckei* in all regions of the genome, including the Z chromosome, with probability ($\varphi_{b1\leftarrow n1}$) around 0.3–0.4 in the noncoding regions and 0.5–0.6 in the coding regions. This is the only substantial introgression estimated in the Z chromosome, with $\varphi_{b1\leftarrow n1} \approx 0.3$, which was still lower than most autosomes. The chromosome 15 inversion region (15b) has a unique history that is much more complex than the rest of the genome; results for 15b are discussed in the next section.

Species divergence time estimates are precise and highly similar across the genome (**Figure S13A**). The posterior means from coding and noncoding loci are strongly correlated, with $\tau_{\text{coding}} \approx x\,\tau_{\text{noncoding}}$ where $x$ varies between 0.5 and 0.6 ($r^2 > 0.95$) in most chromosomal regions (**Figure S14**). We estimate the age of the Melpomene clade ($\tau_r$) to be ~0.020 substitutions per site based on noncoding data and ~0.013 based on coding data. Assuming a constant neutral mutation rate of $2.9\times10^{-9}$ per site per generation (95% CI: $1.3\times10^{-9}$, $1.5\times10^{-9}$) and 4 generations per year (Keightley et al. 2015), we obtain a conservative estimate that the melpomene clade diverged about 1.7 (CI: 0.9, 3.8) million years ago (Ma), which is comparable to a previous estimate of 3.7 (CI: 3.2, 4.3) Ma based on molecular clock dating (Kozak et al. 2015). We estimate present-day and ancestral population sizes ($\theta$) to be of the order of 0.01, largely consistent among chromosomes (**Figure S13B** and **Tables S17** and **18**). For inbred reference individuals (*H. melpomene*, *H. timareta*, *H. numata* and *H. pardalinus*; see **Table S1**), the estimates ($\theta_{\text{Mel}}$, $\theta_{\text{Tim}}$, $\theta_{\text{Num}}$ and $\theta_{\text{Par}}$) tended to be much more variable, with some chromosomes having unusually low estimates in the order of 0.001 or lower. *H. melpomene* has the lowest population size estimate of about 0.002–0.004 on average. This pattern corresponds well to the fluctuating levels of heterozygosity among chromosomes as a result of inbreeding. Adding more individuals should help stabilize the estimates. Inbred individuals should be avoided in population genomic analyses.

#### MSC-M model for pardalinus-hecale and cydno-melpomene clades

In the pardalinus-hecale clade, we find the migration rates ($M_{\text{Par}\rightarrow\text{Ele}}$ and $M_{\text{Ele}\rightarrow\text{Par}}$) between *H. pardalinus* and *H. elevatus* to be high and largely asymmetrical, with $M \approx 2$–4 migrants per generation in one direction and a lower rate ($M < 1$) in the opposite direction (**Figure S15A**), consistent with the introgression probability estimates (**Figure S13A**). However, we observe a label-switching problem in which different runs of model fitting on the same data converged to different posterior peaks that differed in the main direction of gene flow accompanied by a flip in the corresponding population size parameters. This near unidentifiability of $M$ and $\theta$ appears to be partly associated with high migration rates and low genomic distinctiveness between *H. pardalinus* and *H. elevatus*. Since these two species appear to be virtually panmictic and to share effective population size over most of their genome, we also fitted an alternative model to these two species assuming all three populations (two present-day and one ancestral) share the same population size. Under this simpler model, the migration rates are still high ($M$ ~ 1) but less extreme, and the unidentifiability issue largely disappears (**Figure S15B** and **Table S20**). We also detect significant but lesser gene flow involving *H. hecale*, with migration rates $M_{\text{Hec}\to\text{Ele}}$, $M_{\text{Ele}\to\text{Hec}}$, $M_{\text{Hec}\to\text{Par}}$ and $M_{\text{Par}\to\text{Hec}}$ < 0.1 in most chromosomes (**Figure S15A**). Divergence time estimates are more stable among chromosomes and are not affected by the unidentifiability issue. Under this MSC-M model, the divergence time between *H. pardalinus* and *H. elevatus* is $\tau_{\text{ep}} \approx 0.010$ substitutions per site, and the divergence time between *H. hecale* and *H. pardalinus* + *H. elevatus* is $\tau_{\text{eph}} \approx 0.013$ substitutions per site based on noncoding data; coding data led to about half these values (**Figure S15A** and **Table S19**). These estimates are slightly older than those obtained under a larger model assuming pulse introgression in **Figure 2B** are $\tau_{\text{ep}} \approx 0.0070$ and $\tau_{\text{eph}} \approx 0.0076$ using noncoding data ($\tau_{\text{ep}} \approx 0.002$ and $\tau_{\text{eph}} \approx 0.003$ using coding data) (**Tables S17** and **18**).

In the cydno-melpomene clade, estimated migration rates under the MSC-M model are below 0.1, generally much lower than those in the pardalinus-hecale clade, with large variation among chromosomes (**Figure S15C** and **Table S21**). The label-switching problem is rare. Chromosomes with unusually low incoming migration rate estimates tend to be associated with low heterozygosity in inbred individuals (*H. melpomene* and *H. timareta*). For instance, the *H. melpomene* individual had almost no variation in chromosomes 2, 9, 10 and 18 and as a result, the migration rates $M_{\text{Cyd}\to\text{Mel}}$ and $M_{\text{Tim}\to\text{Mel}}$ (and $\theta_{\text{Mel}}$, $\theta_{\text{Tim}}$) are estimated about one or two orders of magnitude lower on those chromosomes compared with other chromosomes (**Table S21**). Unlike the introgression model where introgression is largely from *H. cydno* and *H. timareta* into *H. melpomene*, there is no clear pattern of highly asymmetrical gene flow into *H. melpomene*. In this MSC-M model, we also allow for gene flow between *H. timareta* and *H. cydno*, which is estimated to be $M_{\text{Cyd}\to\text{Tim}} \approx 0.1$ and $M_{\text{Tim}\to\text{Cyd}} \approx 0.04$ migrants per generation in both coding and noncoding loci (**Table S21**). Divergence time estimates are largely comparable across chromosomes. However, the estimates tend to be older than those obtained under the introgression model of **Figure S13A**. We obtained $\tau_{\text{tc}} \approx 0.007$ and $\tau_{\text{tcm}} \approx 0.012$–$0.014$ using coding or noncoding loci under the MSC-M model (**Table S21**) while $\tau_{\text{tc}} \approx 0.0025$ and $\tau_{\text{tcm}} \approx 0.007$ using noncoding data and $\tau_{\text{tc}} \approx 0.001$ and $\tau_{\text{tcm}} \approx 0.0043$ using coding data (**Tables S17** and **S18**).

### References


Dalquen DA, Zhu T, Yang Z. 2017. Maximum likelihood implementation of an isolation-with-migration model for three species. Syst Biol. 66(3):379–398.

Flouri T, Jiao X, Rannala B, Yang Z. 2020. A bayesian implementation of the multispecies coalescent model with introgression for phylogenomic analysis. Mol Biol Evol. 37(4):1211–1223.

Keightley PD, Pinharanda A, Ness RW, Simpson F, Dasmahapatra KK, Mallet J, Davey JW, Jiggins CD. 2015. Estimation of the Spontaneous Mutation Rate in Heliconius melpomene. Mol Biol Evol. 32(1):239–243.

Kozak KM, Wahlberg N, Neild AFE, Dasmahapatra KK, Mallet J, Jiggins CD. 2015. Multilocus species trees show the recent adaptive radiation of the mimetic heliconius butterflies. Syst Biol. 64(3):505–524.

Rosser N, Queste LM, Cama B, Edelman NB, Mann F, Mori Pezo R, Morris J, Segami C, Velado P, Schulz S, et al. 2019. Geographic contrasts between pre- and postzygotic barriers are consistent with reinforcement in Heliconius butterflies. Evolution (N Y). 73(9):1821–1838.

Zhu T, Yang Z. 2012. Maximum likelihood implementation of an isolation-with-migration model with three species for testing speciation with gene flow. Mol Biol Evol. 29(10):3131–3142.
